# Characterization of a Novel Transmembrane Activating STING Agonist using Genetically Humanized Mice

**DOI:** 10.1101/2025.07.14.664814

**Authors:** Nobuyo Mizuno, Jinu Abraham, Kevin Jimenez-Perez, Ian Rose, Laura Springgay, Dylan Boehm, Takeshi Ando, Daniel Streblow, Janine Ward, Shannon Miller, Uddav Pandey, Ahmad Junaid, David Joyner, Roshell Muir, Elias K. Haddad, David Burkhart, Omer Rasheed, Victor R. DeFilippis

## Abstract

STING is a pattern recognition receptor that activates type I interferon and proinflammatory responses in addition to unrelated molecular processes following exposure of DNA to the cytosol. Its pharmacologic stimulation enhances vaccine potency and generates effective antitumor responses but clinical trials evaluating STING agonists have not led to approval for human use. STING activation can occur through ligand engagement of either cytosolic or transmembrane protein domains, processes to which distinct cellular phenotypes are attributed. However, the only transmembrane STING agonist identified is human selective and *in vivo* testing in conventional models is not feasible. Here we describe synthesis of novel STING agonists efficacious against allelic variants of the protein. We also describe genetically humanized STING mice and demonstrate their suitability as a model to evaluate *in vivo* responses following exogenous administration of human-selective agonists. Experiments demonstrate that the lead molecule (termed INI3069) functions through binding to the STING transmembrane region and its comparison with conventional agonists reveals significant differences in molecular and immune effects. INI3069 can also enhance antibody responses to co-administered antigens and antitumor responses. This work both represents the first *in vivo* examination of the effects of transmembrane STING agonism and demonstrates efficacy of a potential novel vaccine adjuvant and oncological therapeutic.

## Introduction

The innate immune response is defined by rapid intracellular reactions to microbe- or danger-associated molecular patterns (i.e. M/DAMPs). These factors are detected by pattern recognition receptors (PRRs) that initiate signal cascades culminating in transcriptomic changes, inflammatory cytokine maturation, and increased antigen uptake and presentation (1, 2). In this way innate immunity both generates tissue states resistant to microbial growth and potentiates adaptive immune responses. <u>St</u>imulator of <u>in</u>terferon <u>g</u>enes (STING) is both PRR reactive to cyclic purine dinucleotides (CDNs) and a signal adaptor for cytosolic DNA-sensing PRRs (3–5). The cellular PRR <u>c</u>yclic <u>G</u>MP-<u>A</u>MP <u>s</u>ynthase (cGAS) is a nucleotidyl transferase that produces 2’-3’ linked GMP-AMP (cGAMP) (3, 6) in response to cytosolic DNA originating from infection (7, 8) as well as transformation-associated processes such as genotoxicity (9), DNA damage responses (10), mitochondrial stress (11), or tumor cell phagocytosis (12–14). Like other PRRs, STING stimulates synthesis of type I interferons (IFN-I), proinflammatory cytokines, co-stimulatory molecules, and antimicrobial effectors (15, 16). However, STING is unusual for reasons related to its diverse molecular functions, stimulus-driven induction, and phenotypic effects (17, 18). First, STING elicits transcriptomes and cytokine profiles (largely through TBK1-mediated activation of transcription factors IRF3 and NF-κB) that are cell type specific yet functionally under characterized *in vivo* (19). Next, STING can activate numerous TBK1-independent processes such as autophagy (20–22), lysosomal biogenesis (23, 24), translational shutoff (25), inflammasome activation (26), and programmed cell death (27). Collectively, these integrate to shape antigen-directed adaptive immunity and the magnitude, proportions, and contexts by which these processes occur vary by stimulus and cell type in ways that require deeper understanding.

The cGAS-STING pathway is crucial to protective innate and adaptive immune processes against infections and cancers [reviewed in (17)]. For instance, mice lacking cGAS, STING, or IRF3 are highly susceptible to viral infection and unable to clear multiple tumor types (12, 28–32). Tumor cell-derived DNA induces IFN-I responses that stimulate dendritic cell (DC) antigen presentation and subsequent tumor cytotoxic CD8^+^ T cells (12). Moreover, STING activation triggers expression of CXCR3 ligands that recruit CD8^+^ cells to the tumor microenvironment (TME) (33). These findings facilitated demonstration that activation of STING using CDNs or small molecules is capable of remarkable tumor clearance and even abscopal responses (34, 35). Additionally, STING-inducing adjuvants enhance the potency of antimicrobial vaccines [reviewed in (36–38)] including generation of both humoral and cell-mediated protection against viral (39–41), bacterial (42, 43), and parasitic (44, 45) infection. Unfortunately, successes observed in mice have not been replicated in clinical trials (46) and the diversity of STING agonist types that have failed as such suggests phenomena other than pharmacokinetics may be relevant. For instance, binding affinity for STING orthologs varies greatly between ligands and this, along with diversity in STING expression between cell types, further influences the immune processes that agonists stimulate (47, 48). Additionally, how TBK1-independent functions vary between agonists and orthologs is largely unexplored especially in relation to the adaptive responses to which they are linked (49). This is particularly evident following engagement of the canonical ligand binding and the transmembrane domains (LBD, TMD) since these agonism modes induce divergent molecular processes (23, 24, 47, 50). Thus, mechanisms connecting the contextual diversity of agonist-induced phenotypes and ensuing adaptive immune responses is lacking yet has crucial implications for clinical targeting of STING. Surprisingly, immune-associated cellular responses elicited by activated STING can vary by chemical structure of the activating ligand (47, 49, 51, 52). As such, the incentive is high to uncover novel STING agonists and to characterize the spectrum of effects they elicit in the context of clinically useful models.

Our group previously identified a molecule that activates human, but not mouse STING (53). We now describe synthesis of more potent analogs obtained through molecular modeling and structure activity relationship (SAR) approaches. To explore the translational applications of these we constructed C57Bl/6 mice in which the endogenous STING coding region was replaced with the human WT allele (huSTING mice). These are capable of reacting *ex vivo* and *in vivo* to human-, but not mouse-selective STING inducers and display enhanced innate and adaptive immune responses when administered antigens are adjuvanted with human STING agonists. We now show that a derivative termed INI3069 can impair tumor growth in these mice and it thus represents a novel human ortholog-selective agonist with anti-tumor properties.

## Results

### Synthesis of novel human-selective STING agonists

G10 is a small molecule that activates human, but not mouse, STING (53). The potency of this molecule was low and as such we used structure-activity relationship and medicinal chemistry to synthesize more effective analogs (**Supplementary Fig. 1**). This led to identification of derivatives designated INI3067, INI3069, INI3070, and INI3071 (**Figure 1A**) that displayed improved peak innate induction and potency as indicated by their ability to stimulate IRF3/IFN-dependent reporter gene expression in human cells that lack the IRF3-terminal adaptor proteins MAVS and TRIF but not STING (**Figure 1B**). Cytotoxicity was also examined and concentration-dependent decreases in cell viability were found to associate with the presence of STING (**Figure 1C**). We next examined induction by the compounds of endogenous mRNAs responsive to IRF3 and IFN-I using semi-quantitative RT-PCR (qPCR). As shown in **Figure 1D**, synthesis of CXCL10, Viperin, and IFIT1 mRNAs was stimulated only in cells that express STING whereas stimuli such as Sendai virus [SeV; (54)] and AV-C (55) inducing MAVS- and TRIF-dependent responses, respectively, are able to activate these genes in STING^-/-^ cells. As an additional control we included the diamidobenzimidazole STING agonist diABZI (56) which also displayed activity only in STING expressing cells. We next used immunoblotting to verify that the canonical STING signaling cascade is activated as expected. As shown in **Figure 1E**, each compound led to phosphorylation of STING, TBK1, and IRF3 but not in cells from which STING was deleted. It is worth noting that INI3069 elicited measurable phosphorylation of TBK1 suggesting that it may trigger residual activation of this protein via an unknown STING-independent mechanism.

**Figure 1.**
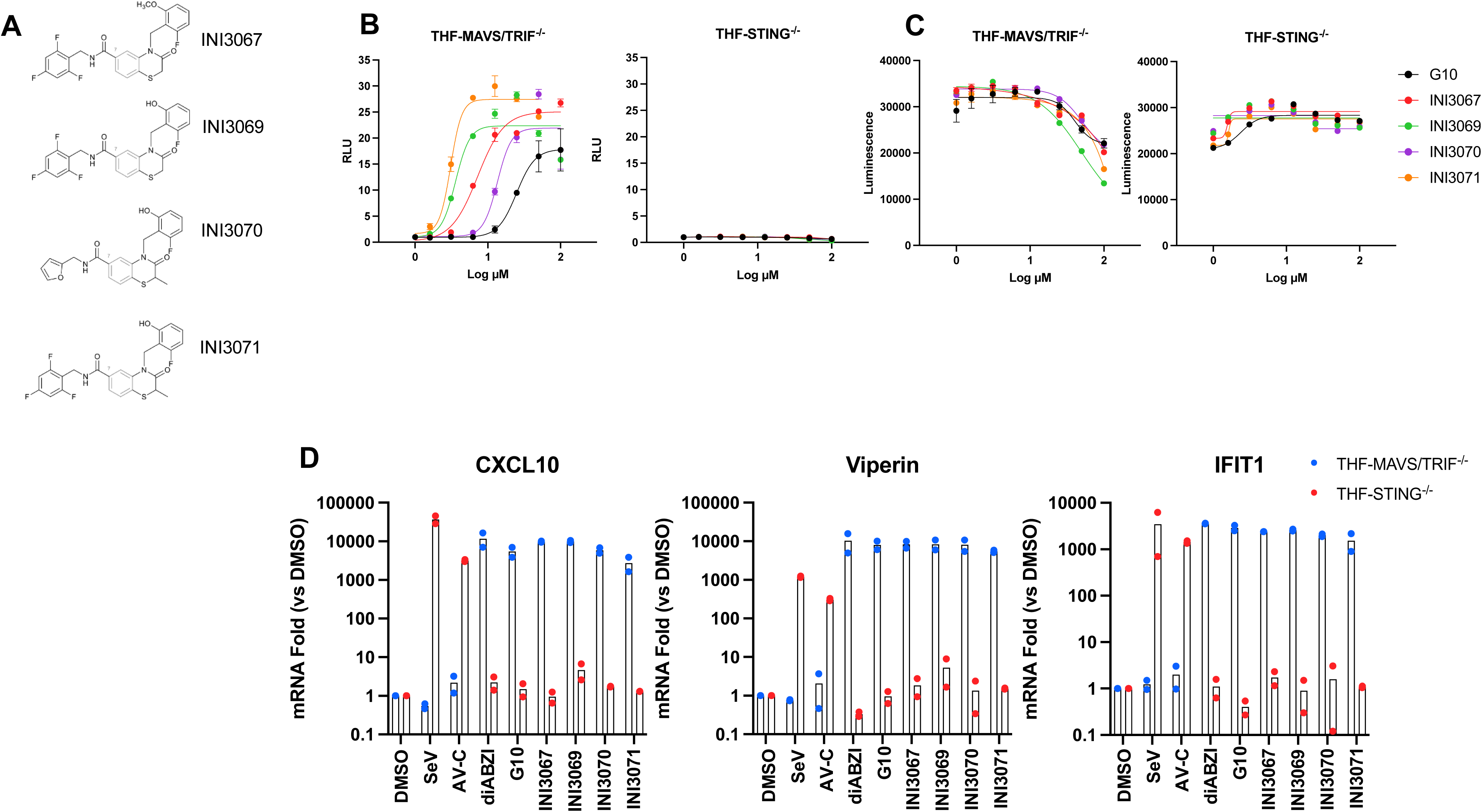
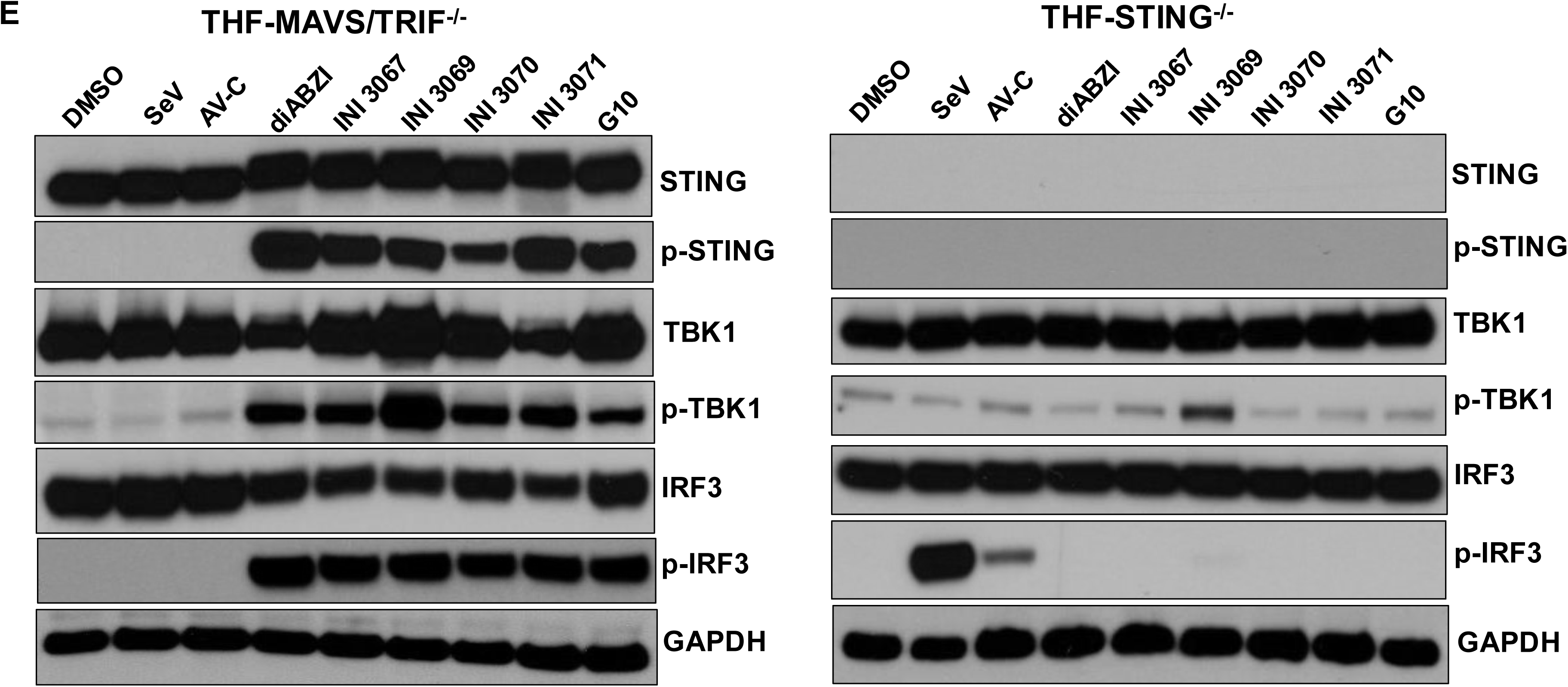
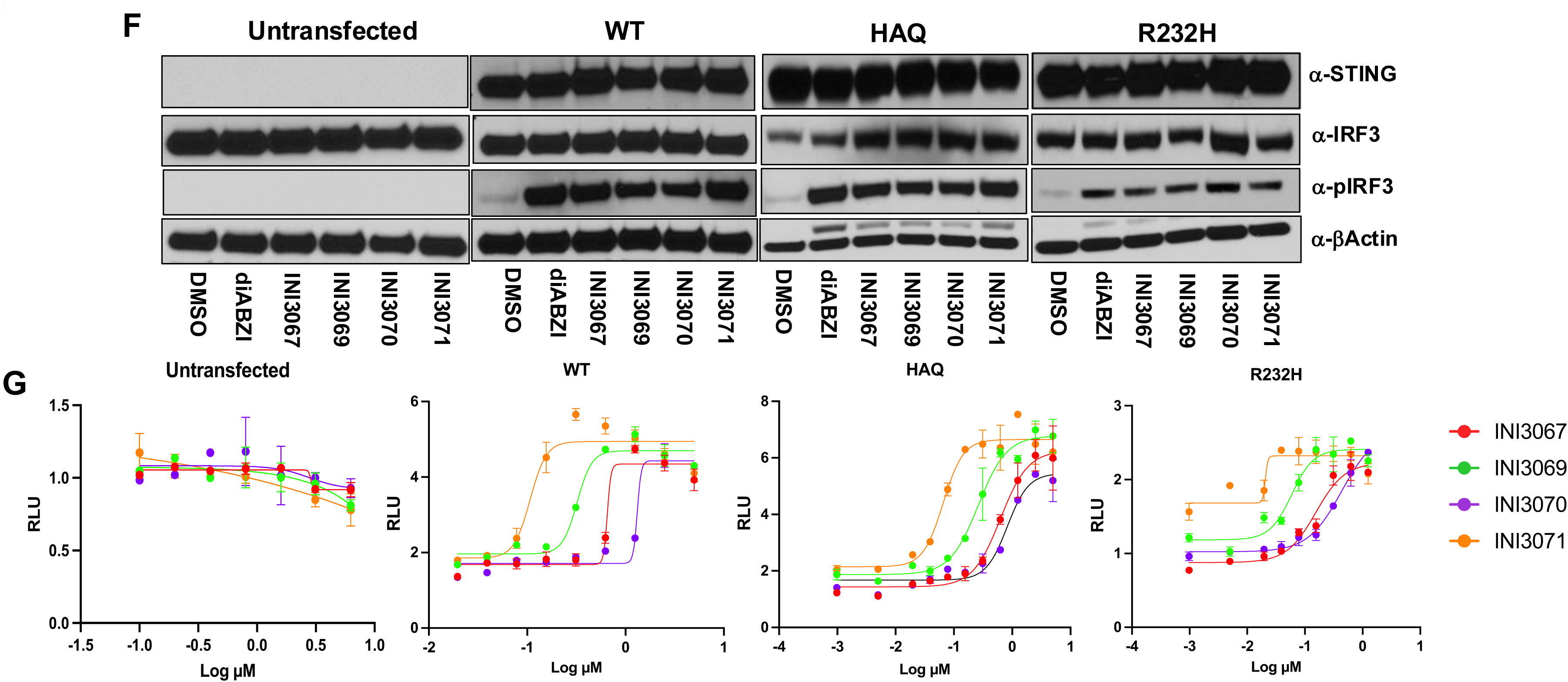
*In vitro* activity of G10 analogs on human cells. **A.** Chemical structures of INI3067, INI3069, INI3070, and INI3071. THF-ISRE cells deficient in TRIF/MAVS or STING were exposed in duplicate overnight to G10 or analogs INI3067, INI3069, INI3070, or INI3071 at indicated concentration; **B.** IRF3/IFN-I-dependent luciferase expression determined by measuring luminescence. Data presented are mean ± SEM relative luminescence units (RLU) determined in relation to dosage matched DMSO-treated cells; **C.** Cell viability was determined using ATP-based Cell Titer Glo assay. Data presented are raw luminescence values; **D.** THF-MAVS/TRIF^-/-^ or THF-STING^-/-^ cells were exposed in duplicate for 6 h to DMSO, 1000 HAU SeV, 100 µM AV-C, 100 nM diABZI, or 100 µM G10, INI3067, INI3069, INI3070, or INI3071. Transcription of mRNAs encoding CXCL10, Viperin, or IFIT1 were then determined using qPCR. Data presented are mean ± SD fold changes of indicated transcript relative to DMSO treated cells; **E.** THF-MAVS/TRIF^-/-^ or THF-STING^-/-^ cells were exposed for 4 h to DMSO, 1000 HAU SeV, 100 µM AV-C, 100 nM diABZI, or 100 µM G10, INI3067, INI3069, INI3070, or INI3071 and immunoblot of whole cell lysates used to determine expression and phosphorylation status of STING, IRF3, and TBK1 with GAPDH as a loading control; **F.** HEK293T cells were left untransfected or transiently transfected for 24 h with vectors expressing the WT, HAQ, or R232H alleles of human STING. Cells were then exposed to DMSO, 100 nM diABZI, or 100 µM INI3067, INI3069, INI3070, or INI3071 for 4 h and immunoblot used to determine expression levels of STING and IRF3 and determine the phosphorylation status of IRF3 with GAPDH as a loading control; **G.** HEK293T cells were stably transfected with indicated STING alleles and transiently transfected with an IRF3/IFN-I-responsive LUC reporter vector and exposed in duplicate overnight to indicated concentrations of INI3067, INI3069, INI3070, or INI3071 or dosage matched DMSO. Data displayed are mean ± SEM RLU.

Whether STING is sufficient for these processes is not demonstrated by these experiments. Moreover, multiple alleles of human STING are described that may display differential reactivity to ligands [reviewed in (57)]. To examine the sufficiency of STING in the response to the analogs and to compare analog efficacy across alleles we stably transduced HEK293T cells [naturally deficient in cGAS and STING (58)] with lentivectors that express one of three common human STING variants [WT (RGRR), HAQ, and R232H (57)] and transiently transfected a luciferase reporter vector responsive to activation of the IRF3/IFN-I-dependent IFIT1 promoter. As shown in **Figure 1F**, in addition to stimulating IRF3 phosphorylation in response to activation of the ectopic STING variants, the compounds exhibited dose dependent IFIT1 reporter activation in the presence, but not absence of each allele. These, along with published data (59), suggest STING is sufficient for G10 analog-mediated IRF3/IFN-I induction *in vitro* in the presence of the three common alleles.

### G10 analogs induce STING- and IRF3-dependent but ATG5-independent antiviral activity

A hallmark of STING activation involves establishment of cellular states that impair viral replication through IFN-I-dependent and autophagic processes (7, 22, 60–64). We therefore examined whether the analogs can elicit antiviral activity *in vitro* and aimed to identify required cellular signaling proteins. For this we pretreated parental THF cells with a dosage range of the four molecules. These were then infected with the (+) RNA Chikungunya virus (CHIKV) engineered to express a nanoluciferase (nLuc) reporter ORF (65). Virus-encoded luminescence levels were measured and compared across treatments. As shown in **Figure 2A**, each compound was capable of stimulating antiviral activity with inhibitory concentrations leading to 50% viral growth inhibition (IC_50_) of 4.895 µM (INI3067), 3.022 µM (INI3069), 5.404 µM (INI3070), and 1.196 µM (INI3071). Since all four analogs potently elicit STING-associated processes, we decided to focus on INI3069 due to the simplicity of its synthesis and low IC_50_. We thus similarly examined the dependence of INI3069 antiviral activity in THF cells lacking MAVS/TRIF, STING, IRF3 (to discern the role of IFN-I pathways) or ATG5 (to discern the role of autophagy). Control stimuli included DMSO (negative) and IFNβ (positive). As shown in **Figure 2B**, IFNβ was able to suppress nLuc signal in all cells as expected since the IFNα receptor (IFNAR) and JAK/STAT pathways are operational. INI3069 suppressed CHIKV in THF-MAVS/TRIF^-/-^ (IC_50_ = 0.6758 µM) and THF-ATG5^-/-^ cells (**Figure 2C**; IC_50_ = 0.5171 µM) but failed to do so in the others. We conclude that INI3069 stimulates antiviral activity in a manner that requires STING and IRF3 but not ATG5 and is thus likely IFN-I-dependent and autophagy-independent.

**Figure 2.**
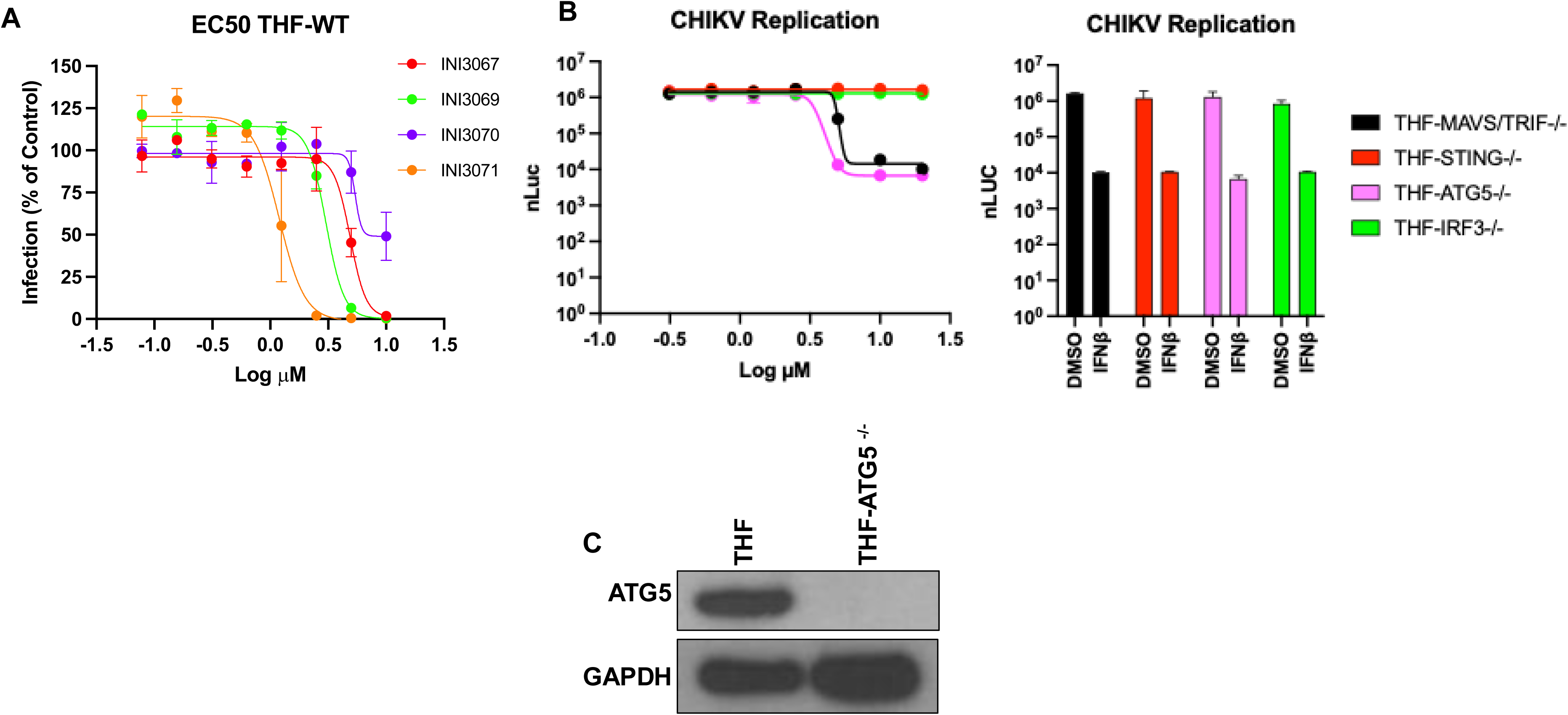
*In vitro* antiviral activity of G10 analogs. **A.** THF cells were treated in duplicate for 6 h with INI3067, INI3069, INI3070, or INI3071 or dosage-matched DMSO and infected with CHIKV-nLuc at MOI = 0.1. At 24 h post infection nLuc expression was measured as luminescence. Data presented are mean ± SEM infection levels indicated as percentage of luminescence signal relative to that generated by DMSO control; **B**. THF cells lacking MAVS/TRIF, STING, IRF3, or ATG5 were treated in duplicate for 6 h with INI3069 at indicated concentration (left), DMSO, or 100U IFNb (right) and infected with CHIKV-nLuc and luminescence measured and displayed as in A; **C.** Immunoblot showing expression of ATG5 in THF or ATG-deficient THF.

### Activity of INI3069 on primary human cells

We next examined whether INI3069 can trigger innate phenotypes on primary human cells. For this we obtained peripheral blood mononuclear cells (PBMC) from six healthy donors and first asked whether INI3069 could induce maturation of DCs from these. For this we exposed the PBMCs *ex vivo* to INI3069 at two concentrations or dose-matched DMSO. Flow cytometry was then used to quantify surface expression of maturation markers CD86 and CD83. Significant increases were detectable for both markers at both concentrations (**Figure 3A**). We next used a multiplex immunoassay to quantify secretion of proinflammatory cyto/chemokines following similar treatment. As shown in **Figure 3B**, significant induction of eighteen cyto/chemokines was detected following treatment with either one or both concentrations examined. Based on this we conclude that INI3069 activates innate processes in primary human cells and thus may be suitable for clinical applications.

**Figure 3.**
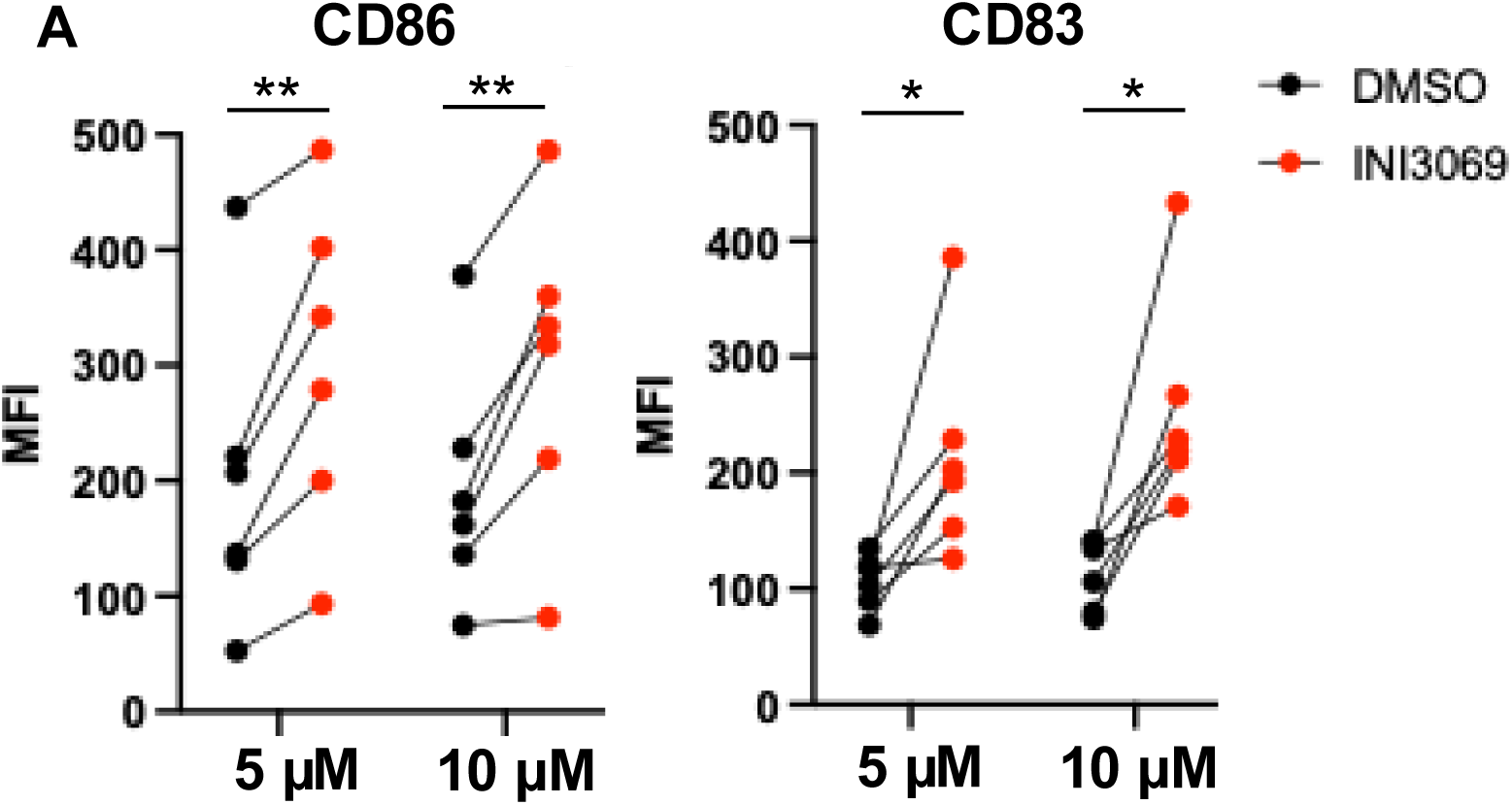
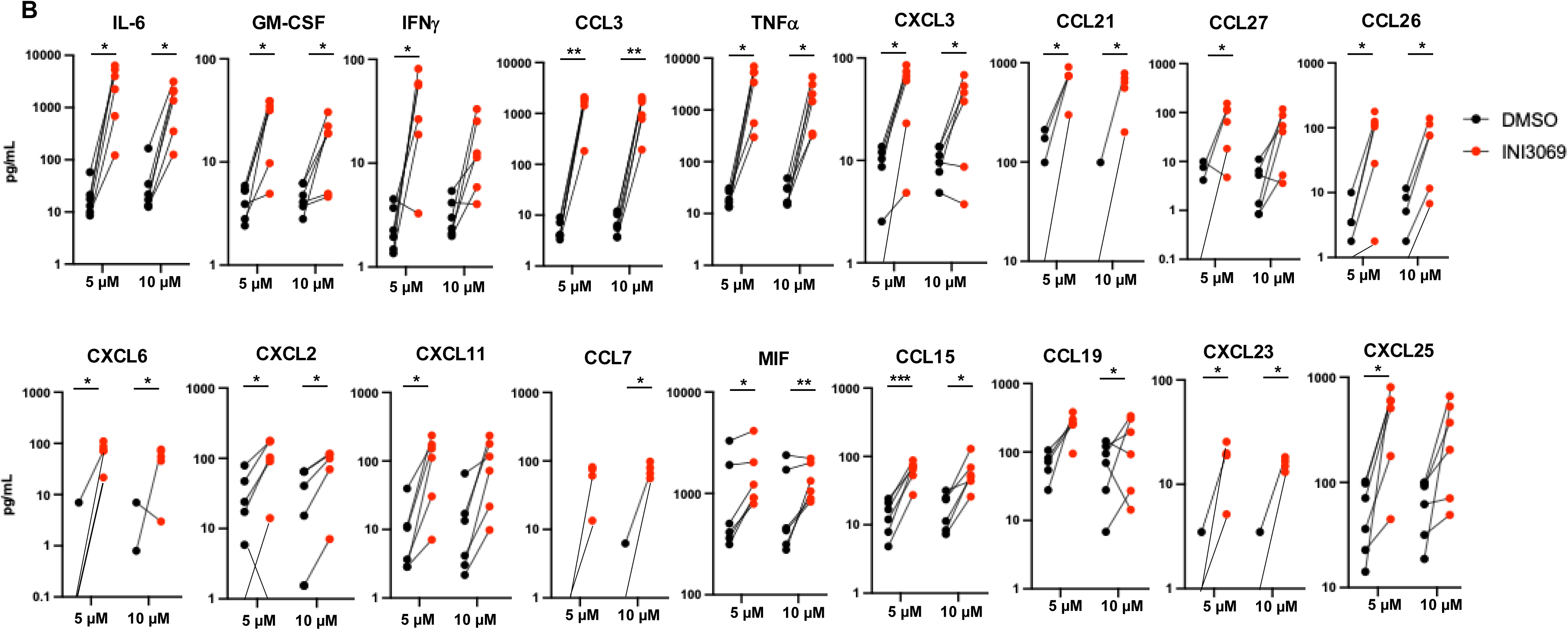
Reactivity of primary human cells to INI3069. PBMCs were obtained from six healthy human donors and grown *ex vivo*. **A.** Immature DCs were obtained from PBMC and exposed 24 h to INI3069 at 5 or 10 µM or dosage matched DMSO vehicle in RPMI + 10% FCS. Flow cytometry was then used to quantify expression of CD83 or CD83. Data presented are geometric mean fluorescence intensities (MFI) of cell populations from individual donors; **B.** Cells were exposed to 5 or 10 µM INI3069 or a dosage matched concentration of DMSO vehicle for 24 h. Media was harvested and secreted cytokines measured using Luminex assay. Data presented are pg/mL of indicated cytokine for cell cultures from individual donors. Statistical significance was determined using paired sample T-test (*p < 0.05; **p < 0.01, ***p < 0.001).

### INI3069 stimulates STING activation by engaging the transmembrane domain

STING includes a four-pass transmembrane domain (TMD) that anchors it to the ER as well as a cytosolic region that contains a CDN ligand binding domain (LBD) and C-terminal tail required for recruitment of TBK1 and IRF3 (66). While ligands that engage the LBD (67) such as diABZI and cGAMP are most common, a human selective agonist termed C53 has been described that activates via the TMD (68). Importantly, this directs the cellular processes stimulated by activated STING (23, 24, 47, 69). To identify the domains required for binding of INI3069 we first purified the STING C-terminal cytosolic domain (CTD) and used a fluorescence based thermal shift assay (70). As shown in **Figure 4A**, combining STING-CTD with cGAMP led to a substantial change in thermal stability as indicated by the shift in fluorescence emission with increasing temperature. However, INI3069 did not affect thermal stability as emission resembled that observed by STING alone. To determine whether INI3069 activates STING by engaging the TMD we constructed plasmids that express versions of the protein that contain previously described point mutations in targeted domains known to be essential to C53 (67, 71). These, along with a control vector expressing WT STING, were then transfected into HEK293T cells and treated with DMSO, diABZI, C53, or INI3069. Whole cell lysates were then harvested and immunoblot used to determine the phosphorylation status of IRF3 as an indicator of active STING signaling. As shown in **Figure 4B**, all stimuli triggered phosphorylation of IRF3 in the presence of WT STING. However, expression of TMD mutants L30A or Y104A abrogated the ability of C53 and INI3069, but not diABZI, to induce this response. Based on these observations we conclude that INI3069 activation of STING requires the TMD, but not LBD, region, a finding that can be used in future SAR modeling studies to develop improved analogs.

**Figure 4.**
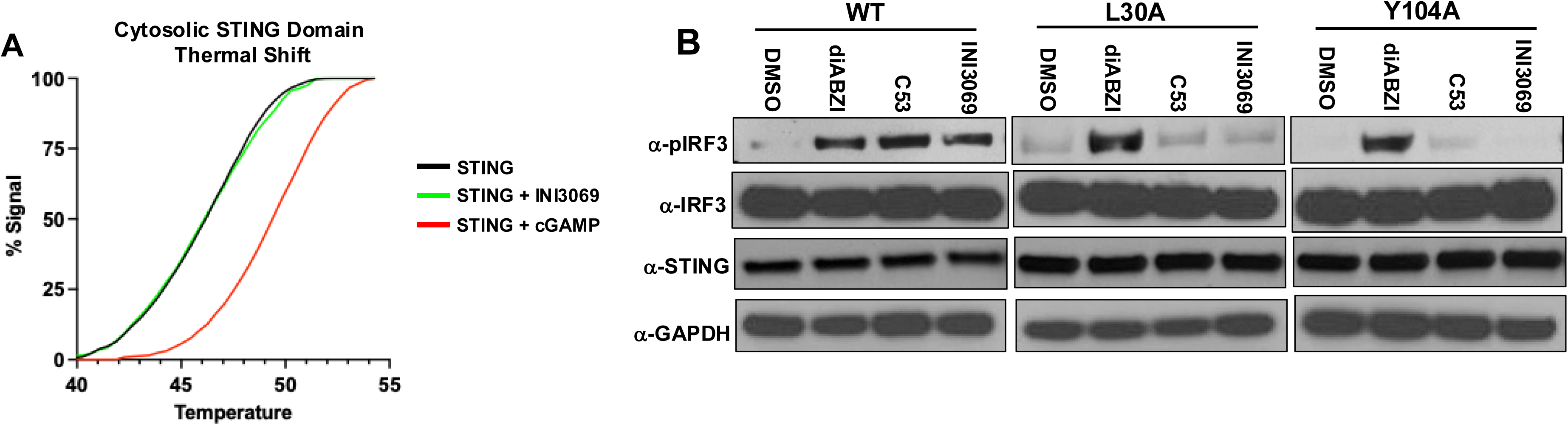
INI3069-mediated activation requires amino acids in the STING transmembrane region and fails to activate TFEB. **A.** 6xHIS-tagged cytosolic domain of STING was synthesized in BL21 bacteria and purified. It was then untreated or mixed with 50 µM INI3069 or 100 µM cGAMP in the presence of SYPRO orange. Fluorescence was measured over the indicated temperature gradient. Data displayed are fluorescence signal for indicated treatments at indicated temperatures; **B.** HEK293T cells were transiently transfected overnight with vectors encoding the WT human STING allele or STING coding regions containing alanine mutations in the TMD (L30A, Y104A) as indicated. Cells were then exposed for 4 h to 0.5% DMSO, 10 nM diABZI, 100 µM C53, or 6.25 µM INI3069. SDS-PAGE was then used on whole cell lysates and stained for phosphorylated IRF3 or STING; **C.** THF-MAVS/TRIF-/- cells were treated 3 h with DMSO, cGAMP, C53, or INI3069. SDS-PAGE was then used on whole cell lysates and stained for phosphorylated TFEB, total TFEB, phosphorylated IRF3, total IRF3, or b-Actin.

### INI3069 fails to activate TFEB or STING-mediated autophagy

STING oligomerization following LDB-mediated activation can form a proton channel independently of TBK1 that raises Golgi and diminishes cytosolic pH (47). This, in turn, leads to GABARAP lipidation and inhibition of mTORC1, a kinase that maintains phosphorylation of the transcription factor TFEB (23, 24). Dephosphorylated TFEB translocates to the nucleus where it upregulates genes necessary for lysosomal biogenesis, a process that enhances direct antigen presentation (72). C53 binding to the TMD is known to block proton leakage and in this way fails to activate TFEB (23, 24, 50, 69) and we therefore asked if INI3069 behaves similarly. As shown in **Figure 5A**, TFEB in THF-MAVS/TRIF^-/-^ cells is dephosphorylated in response to treatment with diABZI as expected. However, TFEB remained phosphorylated following treatment with C53 or INI3069 despite their ability to induce IRF3 phosphorylation suggesting that INI3069 and C53 fail to stimulate the TBK1-independent effects observed for diABZI.

**Figure 5.**
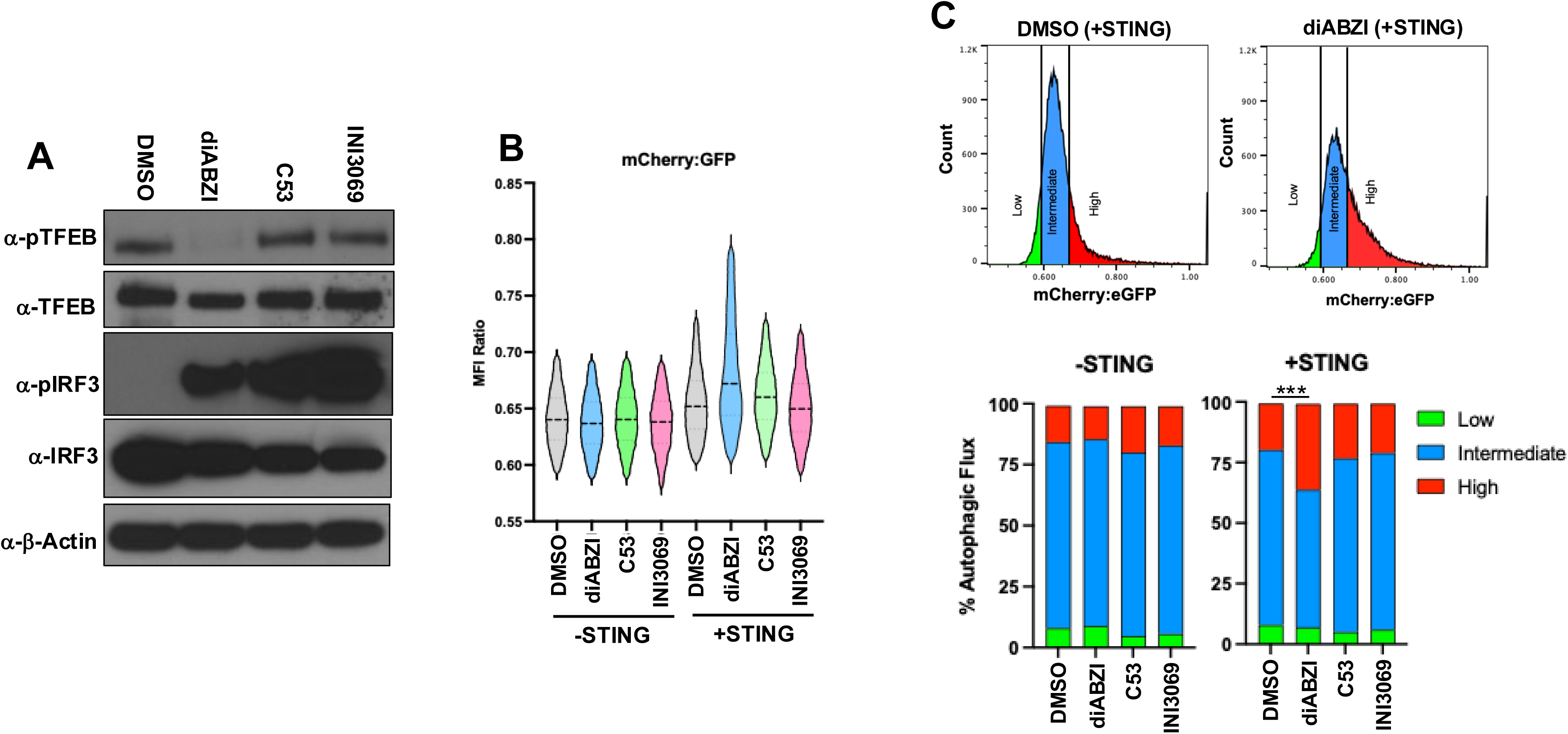
INI3069 fails to activate TFEB and autophagy in human cells. **A.** THF-MAVS/TRIF^-/-^ cells were treated 3 h with DMSO, cGAMP, C53, or INI3069. SDS-PAGE was then used on whole cell lysates and stained for phosphorylated TFEB, total TFEB, phosphorylated IRF3, total IRF3, or b-Actin; **B** and **C.** HEK293T cells were stably transduced with GFP-mCherry-LC3 and two single cell clones transfected for 24 h with a plasmid encoding WT human STING or left untransfected. They were then exposed for 4 h to DMSO, 1 µM diABZI, 10 µM C53, or 10 µM INI3069 and flow cytometry used to quantify intracellular expression of mCherry and GFP. **B.** Summary of mCherry:GFP MFI ratio for individual cells (excluding the top and bottom 10% of observations); **C.** Representative designations of degree of autophagic flux as determined by mCherry:GFP ratio between treatments. Strategy for designating low, intermediate and high autophagic flux based on mCherry:GFP MFI ratio (top). Data presented are proportion of cells displaying indicated level of autophagic flux following indicated treatment in the presence and absence of transfected STING (bottom). Experiments were performed in duplicate for both clonal cell lines and one way ANOVA with Dunnet’s correction for multiple comparisons used to determine statistical significance of cell populations designated as displaying high autophagic flux (***p < 0.001).

STING-mediated proton leakage is also linked to induction of noncanonical autophagy through sensing of vesicle deacdification by vaculoar ATPase (V-ATPase) which recruits ATG16L1 and catalyzes LC3 lipidation (22, 73). To address whether this process is blocked by INI3069 we constructed HEK293T cell clones that stably express a protein chimera comprised of mCherry, GFP, and LC3 to quantify the fusion of autophagosomes and lysosomes (autophagic flux) (74). Due to the sensitivity of GFP fluorescence to lysosomal degradation, flow cytometry is used to determine the relative signal of mCherry and GFP, the ratio of which increases with autophagic flux. These cells were transfected with a plasmid expressing human STING or left untransfected and exposed to DMSO, diABZI, C53, or INI3069. As shown in **Figure 5B**, no stimuli led to significant changes in mCherry:GFP MFI ratios in the absence of STING. However, in the presence of STING, the fraction of cells displaying a high autophagic response increased significantly following exposure to diABZI but not C53 or INI3069 (**Figure 5C**). These results indicate that while INI3069 activates STING-mediated IRF3 phosphorylation, it fails to activate autophagy, a finding consistent with the lack of impact of ATG5 on INI3069-mediated antiviral responses and its engagement of the STING TMD and likely blockade of proton leakage.

Replacement of the endogenous coding region for STING with the human ortholog yields viable animals and functional *in vivo* expression of the ectopic ORF.

G10 analogs fail to activate murine STING and, as such, examining their *in vivo* efficacy in conventional models is not feasible. Since STING-deficient mice are healthy (75) and previous work has demonstrated the capacity of ectopic human STING to activate innate signaling in mouse cells (76, 77) we hypothesized that replacement of the mouse coding region with an open reading frame (ORF) that encodes the human ortholog would lead to generation of viable animals responsive at molecular and immunologic levels to human selective STING inducers. To accomplish this, embryonic cell homologous recombination was performed by electroporation of a vector containing the sequence for the coding portion of human *Tmem173* as illustrated in **Figure 6A**. Presence of the transgenic insert was confirmed by PCR using primer that bind internally to the engineered locus as well as externally in the targeted chromosomal region isolated from animal tissues (**Figure 6A, 6B**). Animals were bred to homozygosity and absolute levels of human STING mRNA transcripts compared across tissue types to levels of endogenous STING mRNA from parental C57Bl/6 mice using qPCR. As shown in **Figure 6C**, significant differences in the respective mRNA levels were only detected between parental and huSTING mice in heart muscle (higher in huSTING mice) and bone marrow derived dendritic cells (BMDC; lower in huSTING mice). These data suggest that across cellular environments homeostatic regulation of transcription and mRNA turnover of the huSTING transgene largely align with that observed for endogenous wild type allele.

**Figure 6.**
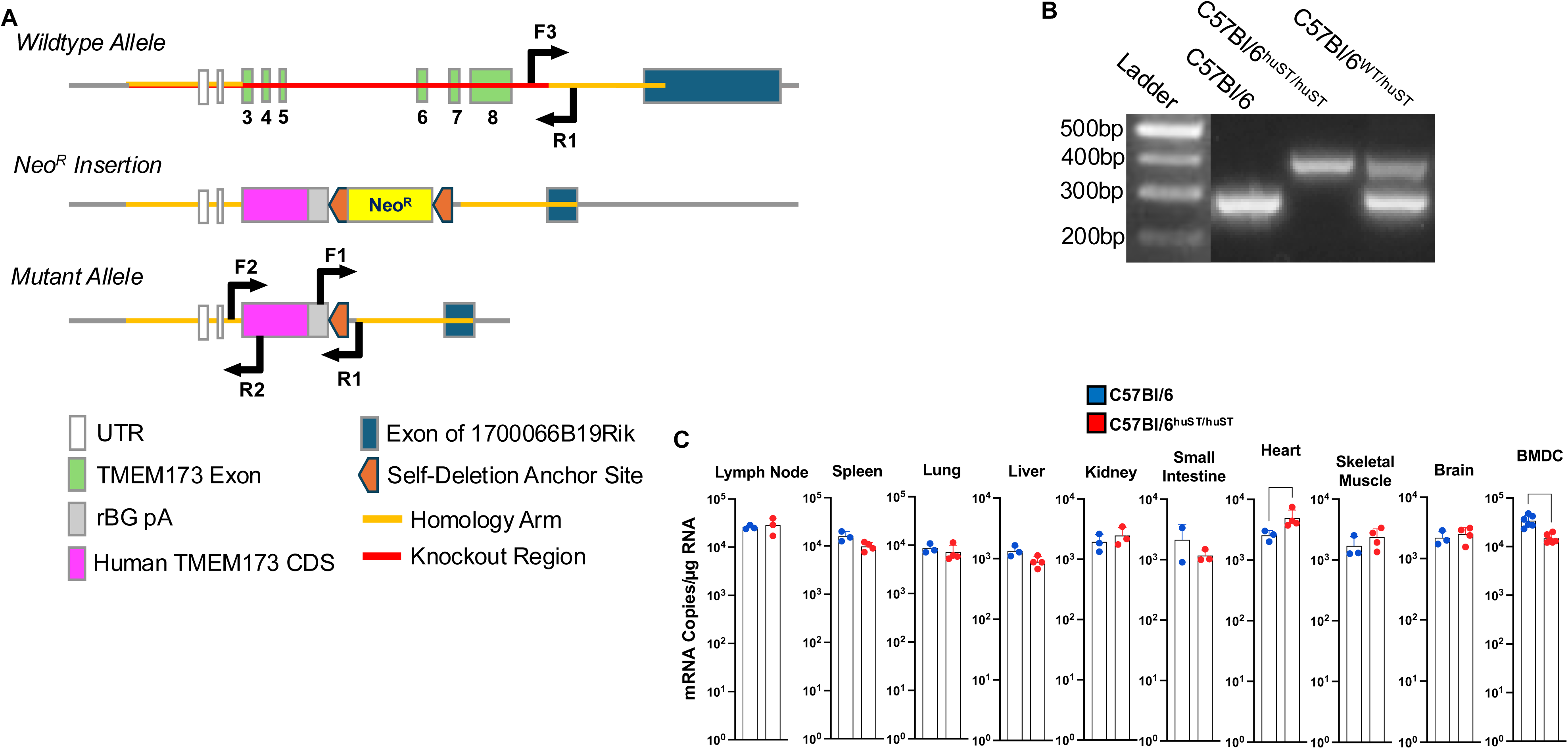
Genomic knockin of coding region for WT allele of human STING. **A.** Schematic illustration of human STING targeting construct, knock in strategy and location of diagnostic PCR primer pairs; **B.** PCR using genomic DNA from indicated parental, heterozygotic, and homozygotic animal strains; **C.** Quantitative RT-PCR using template RNA isolated from indicated tissue and mouse strain. Values are individual (circles) and average ± SD copy numbers of mRNA per μg total RNA.

### Comparative molecular function of transgenic huSTING in response to activating ligands

Regulation of STING is complex, involving multifaceted feedback mechanisms (78). Moreover, whether the huSTING protein signals in a biologically appropriate manner in the context of murine cells, especially following ligand engagement, is crucial to characterizing *in vivo* systemic STING-mediated phenotypes. To characterize this, we began by examining activation of transgenic STING-mediated molecular signaling. Murine embryonic fibroblasts (MEF) were prepared from C57Bl/6, huSTING, and STING-deficient (Goldenticket; STING^Gt/Gt^) mice (79). These were then exposed to SeV as a STING-independent control and agonists of STING that are either species selective or cross activating and harvested for immunoblot. As shown in **Figure 7A**, SeV induced phosphorylation of TBK1 and IRF3 in all cells. STING, TBK1, and IRF3 from all but STING^Gt/Gt^ cell types was phosphorylated as expected following treatment with diABZI. Small molecules that induce species selective activation of STING proteins include DMXAA (mouse) (80) and INI3069. As shown in **Figure 7A**, the compounds only activated phosphorylation of STING, TBK1, and IRF3 in a manner consistent with their known species specificity and none triggered these responses in STING^Gt/Gt^ MEFs. In line with observations made on STING^-/-^ human cells (**Figure 1D**), INI3069 was also able to stimulate residual phosphorylation of TBK1 in C57Bl/6 and STING^Gt/Gt^ MEFs suggesting off target or noncanonical activation of this response. Interestingly, INI3069 also led to detectable phosphorylation of IRF3 in C57Bl/6 but not STING^Gt/Gt^ MEFs and thus a noncanonical STING-dependent mechanism may be involved.

**Figure 7.**
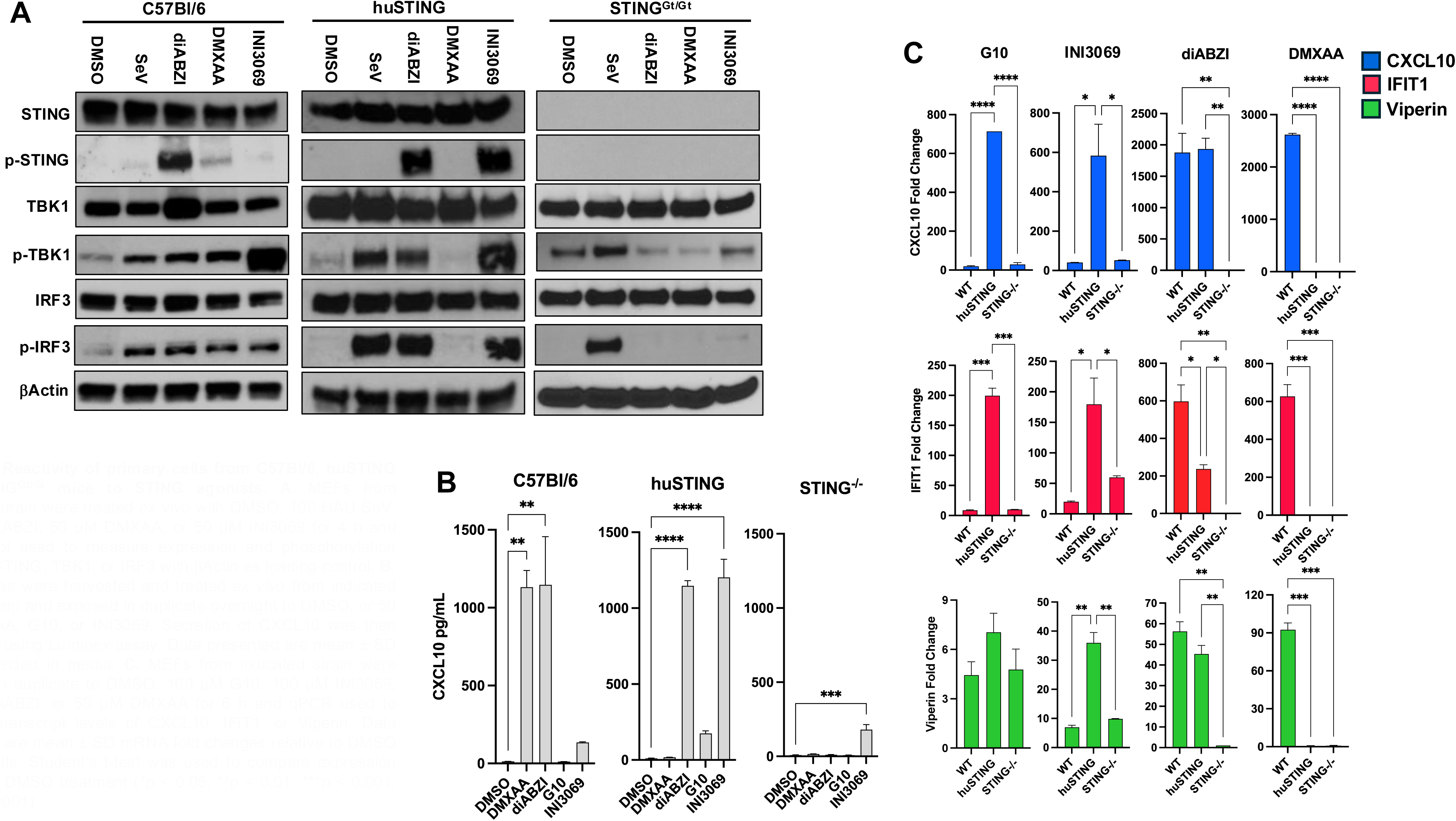
Reactivity of primary cells from C57Bl/6, huSTING and STING^Gt/Gt^ mice to STING agonists. **A.** MEFs from indicated strain were treated *ex vivo* with DMSO, 100 HAU SeV, 100 nM diABZI, 50 µM DMXAA, or 50 µM INI3069 for 4 h and immunoblot used to measure expression and phosphorylation status of STING, TBK1, or IRF3 with bActin as loading control; **B.** Splenocytes were harvested and treated *ex vivo* from indicated mouse strain and exposed in duplicate overnight to DMSO, or 50 µM DMXAA, G10, or INI3069. Secretion of CXCL10 was then measured using Luminex assay. Data presented are mean ± SD levels detected in media; **C.** MEFs from indicated strain were exposed in duplicate to DMSO, 100 µM G10, 100 µM INI3069, 100 nM diABZI, or 50 µM DMXAA for 6 h and qPCR used to measure transcript levels of CXCL10, IFIT1, or Viperin. Data presented are mean ± SD mRNA fold changes relative to DMSO treated cells. Student’s t-test was used to compare expression relative to DMSO treatment (*p < 0.05, **p < 0.01, ***p < 0.001, ****p < 0.0001).

We next examined whether these agonists can induce convergent transcriptional outcomes in primary immune cells. For this we isolated splenocytes from C57Bl/6, huSTING, and STING^Gt/Gt^ mice. Secretion of the chemokine CXCL10 was first examined using Luminex assay. As shown in **Figure 7B**, diABZI treatment led to secretion of CXCL10 in WT and huSTING but not STING^Gt/Gt^ splenocytes whereas DMXAA only induced CXCL10 secretion in splenocytes from WT mice. While G10 induced detectable CXCL10 secretion in huSTING (but not WT or STING^Gt/Gt^) splenocytes, INI3069 induced high secretion in huSTING and lower, but measurable, secretion in WT and STING^Gt/Gt^ splenocytes consistent with observed residual induction of STING-independent innate activity. qPCR was next used to more precisely validate the induction of IRF3/IFN-stimulated genes (ISGs; CXCL10, Viperin, and IFIT1) following *ex vivo* stimulation with the STING inducers. As shown in **Figure 7C**, diABZI led to ISG induction in wild type and huSTING, but not STING^Gt/Gt^ splenocytes. As expected, DMXAA only induced ISG expression in splenocytes from wild type mice. G10 and INI3069 induced substantial ISG expression in splenocytes from huSTING mice while INI3069 induced detectable IFIT1 and Viperin induction in C57Bl/6 and STING^Gt/Gt^ splenocytes. Collectively, these data indicate that molecular function of exogenously stimulated transgenic human STING in the context of murine cells reacts to ligand engagement in a manner that resembles that seen in cells from wild type mice.

### Transcriptomes of dendritic cells from huSTING and C57Bl/6 mice in response to innate stimuli

Data presented in **Figure 7** suggest that agonist-mediated STING signaling by transgenic huSTING resembles that seen via endogenous murine STING. However, myriad mRNAs are known to be regulated in response to STING activation, perhaps even in the absence of ligand engagement (81). Moreover, since immunologically critical BMDC appear to synthesize huSTING mRNA at lower levels than their WT counterparts (**Figure 6B**), we examined the implications of both the ortholog dissimilarity and differential expression levels in C57Bl/6 and huSTING mice on agonist-induced activity in BMDC treated *ex vivo*. For this, BMDC from wild type and huSTING were cultured and treated with cGAMP and diABZI as well LPS as a STING-independent IRF3-activating control stimulus. Total RNA was then harvested at 6 h post treatment, an early time point chosen to diminish the impact of autocrine/paracrine effects of secreted cytokines (82). Hybridization array was then used to quantify levels of >20,000 individual transcripts. With the exception of *Tmem173* (STING; not shown), which expectedly does not cross-hybridize to the mouse array, no transcripts were identified as significantly differentially expressed in control DMSO-treated BMDC between huSTING and WT cells (**Supplemental Table 1**). Furthermore, principal component analysis (PCA) of absolute mRNA levels revealed that transcriptomes segregate predominantly by stimulus rather than by mouse strain (**Figure 8A**). Fold changes of transcripts induced by each stimulus were also highly correlated between wild type and huSTING cells (**Figure 8B**). Likewise, predicted regulation of biologically characterized pathways (**Supplemental Table 2**) was also highly similar between the strains for the different STING agonists as illustrated in **Figure 8C**. These results indicate that cells from huSTING and wild type mice do not react in an overtly transcriptionally dissimilar manner to exogenous activators of STING or STING-independent stimuli. In addition, they also illustrate that STING-dependent transcription can vary in relation to activating ligand (**Figure 6A**).

**Figure 8.**
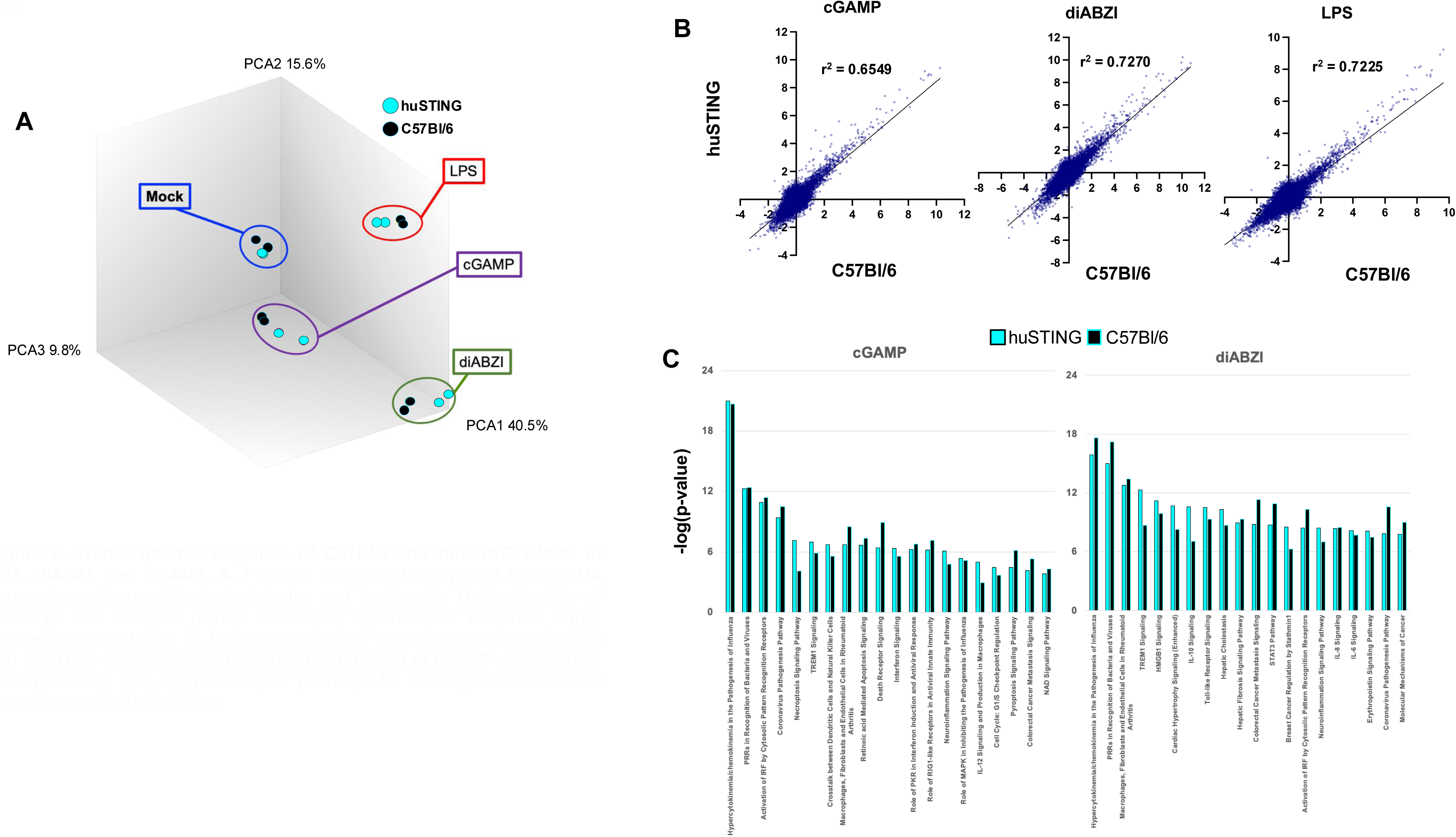
Transcriptomic response of C57Bl/6 and huSTING BMDC to LPS, diABZI, and cGAMP. **A.** Principal component analysis of total mRNA expression patterns of indicated strains and treatments; **B.** Scatterplots of transcript fold changes relative to DMSO treatment for indicated stimuli between mouse strains; **C.** Pathway analysis of transcriptomes from C57Bl/6 and huSTING BMDC following treatment with diABZI or cGAMP as indicated. Data presented are –log(p-values) of pathways for which positive Z scores were observed in both strains.

### STING agonist-induced activity on huSTING BMDC

Antigen-directed adaptive immune responses require activation and maturation of antigen presenting cells (APC) such as DCs in response to innate stimuli. These are particularly relevant to humoral and cell mediated responses for which STING activation has been shown to be a potentiator (83). We therefore examined whether immature BMDCs from huSTING mice respond as such through the expression of conventional markers of APC maturation following exposure to diABZI and INI3069. Immature BMDCs from wild type and huSTING mice were harvested and treated as described above. Flow cytometry was then used to measure surface expression of maturation markers CD40, CD80, and CD86. As shown in **Figure 9**, diABZI was able to significantly increase expression of all three markers in BMDC from both mouse strains. Alternatively, INI3069 only induced maturation in cells from huSTING mice, consistent with results described above. These data indicate species-predicted, STING-dependent innate immune responsiveness at molecular and transcriptional levels in cells from huSTING mice and suggest that the observed decrease in the expression STING mRNA in BMDC relative to WT mice (**Figure 5B**) has no substantial impact.

**Figure 9.**
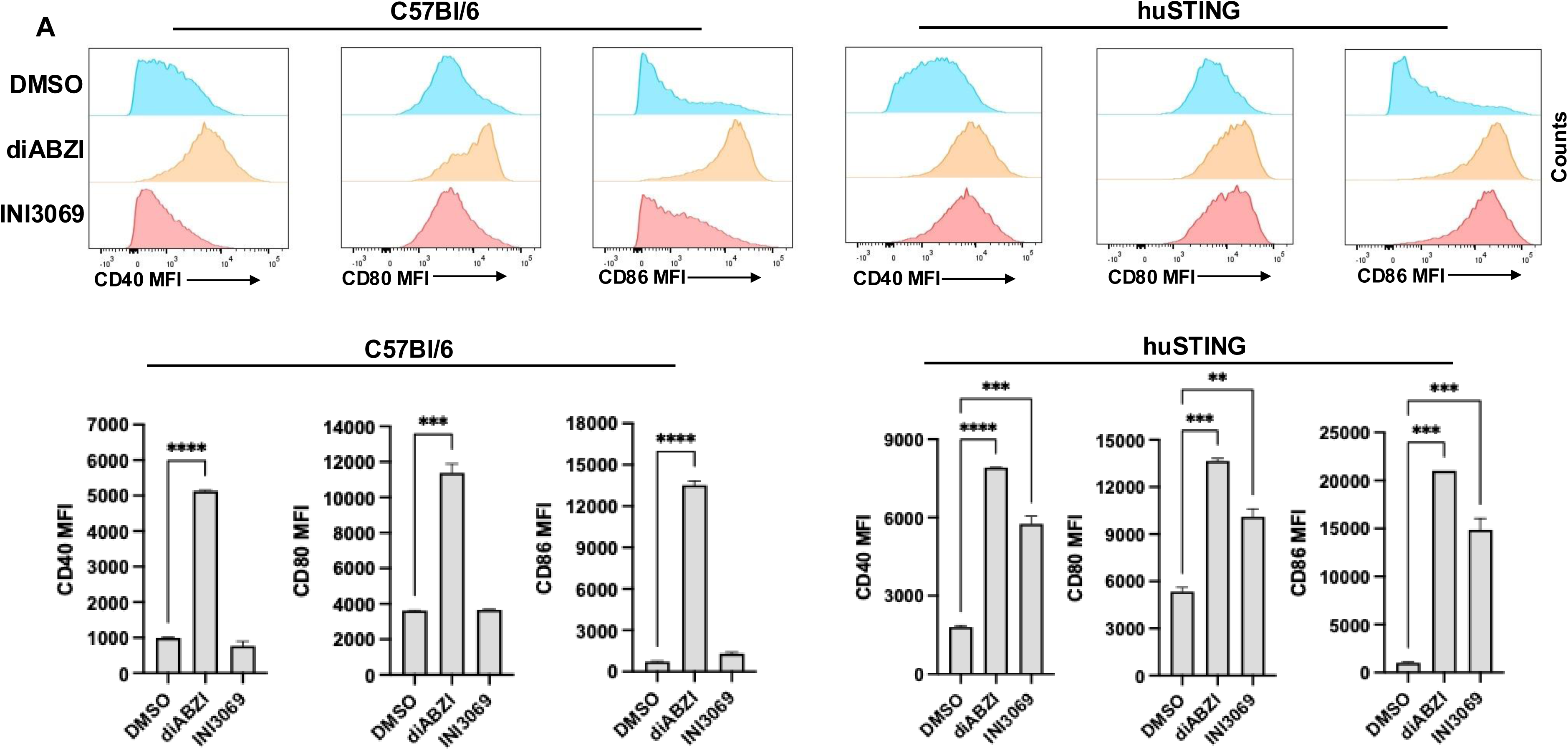
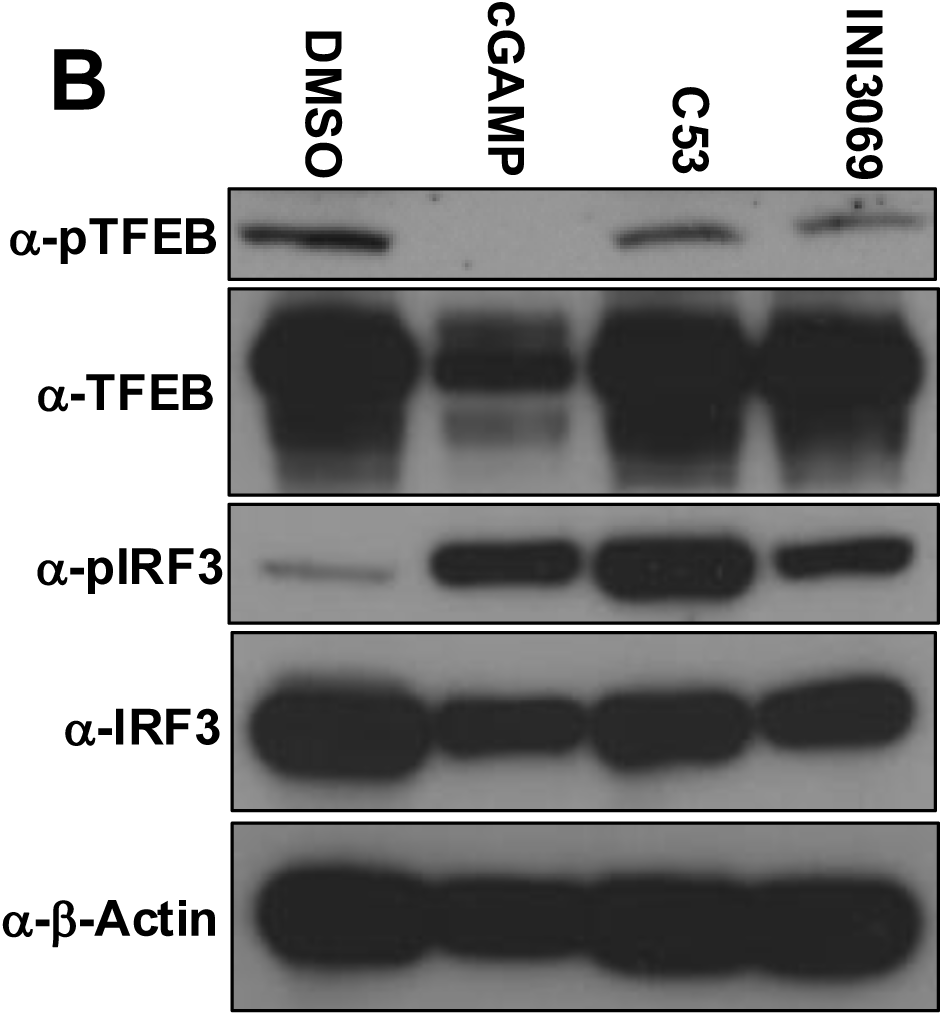
BMDC from C57Bl/6 and huSTING mice respond to INI3069. **A.** Immature BMDC were generated from bone marrow of indicated strain as described in text and treated in duplicate for 24 h with DMSO, 10 nM diABZI, or 10 µM INI3069. Flow cytometry was then used to measure surface expression of CD40, CD80, and CD86. Data presented are sample flow cytometry plots for individual samples (top) and mean ± SD MFI for indicated markers (bottom). One way ANOVA was used to determine significance (**p < 0.01, ***p < 0.001, ****p < 0.0001); **B.** BMDC from huSTING mice were treated 3 h with DMSO, cGAMP, C53, or INI3069. SDS-PAGE was then used on whole cell lysates and stained for phosphorylated TFEB, total TFEB, phosphorylated IRF3, total IRF3, or b-Actin.

Cells from huSTING mice react expectedly to activating ligands by generating canonical TBK1/IRF3-dependent and innate cell maturation responses. However, the TBK1-independent proton channel function of STING also elicits intracellular processes that likely affect immune activity although this has not been examined *in vivo*. Whether the protein behaves similarly in the context of mouse innate cells is important for both establishing the suitability of these animals as a model and to validate the activity of INI3069 observed in human cells. We therefore examined STING agonist-mediated dephosphorylation of TFEB in BMDC. As shown in **Figure 9C**, dephosphorylation of TFEB following treatment of BMDC harvested from huSTING mice with cGAMP occurs as expected. However, while C53 and INI3069 are active in these cells as indicated by their ability to induce IRF3 phosphorylation, they fail to stimulate dephosphorylation of TFEB as was observed in human cells (**Figure 7C**). These results suggest that INI3069 resembles C53 in that it activates STING-dependent IRF3 phosphorylation but not TFEB. Moreover, huSTING behaves similarly when expressed as a transgene in mouse cells to endogenous STING in human cells when activated by LBD-versus TMD-binding agonists and thus strengthening this model.

### INI3069-associated enhancement of antigen-reactive humoral responses in huSTING mice

Agonists of STING can augment the immunogenicity of co-administered protein antigens that manifests as an increase in titers of antigen-reactive serum antibodies [reviewed in (37, 84)]. This has been shown to elicit anti-tumor immune effects when STING agonists are admixed with tumor-expressing antigens including ectopic ovalbumin (OVA) (85–87). We therefore examined whether subcutaneous (SQ) injection of OVA paired with INI3069, CDA, diABZI, or Alum as control adjuvants using a prime-boost schedule could elicit OVA-specific IgG in huSTING and WT mice. As shown in **Figure 10A** and **10B** diABZI, a CDA derivative modified for *in vivo* use (88), and Alum expectedly led to significant increases in titers of OVA-reactive total IgG relative to antigen adjuvanted with formulation vehicle alone in both huSTING and WT mice. INI3069 led to significantly enhanced IgG titers in huSTING but not WT mice, a finding consistent with the compound’s capacity for ortholog-specific innate induction and proper functioning of the protein. T helper (T_H_) polarization can also be affected by adjuvant class with STING agonists having demonstrated a range of T_H_1, T_H_2, and T_H_17 responses largely in relation route of administration and chemical structure [reviewed in (37)]. As an indicator of T_H_ polarization induced by INI3069 relative to control agonists we quantified OVA-reactive IgG1 and IgG2c isotypes. As shown in **Figure 10C**, all adjuvants led to significant and similar induction of IgG1 isotypes. However, Alum and INI3069 elicited substantially lower IgG2c isotype levels than did CDA or diABZI. These results suggest a more balanced T_H_1/ T_H_2 polarization for CDA and diABZI and higher T_H_2 polarization for Alum and INI3069 underscoring key immunological differences between STING agonist classes. This may be functionally linked to the different regions of STING binding and TFEB activation for these agonist classes but will require additional study to elucidate.

**Figure 10.**
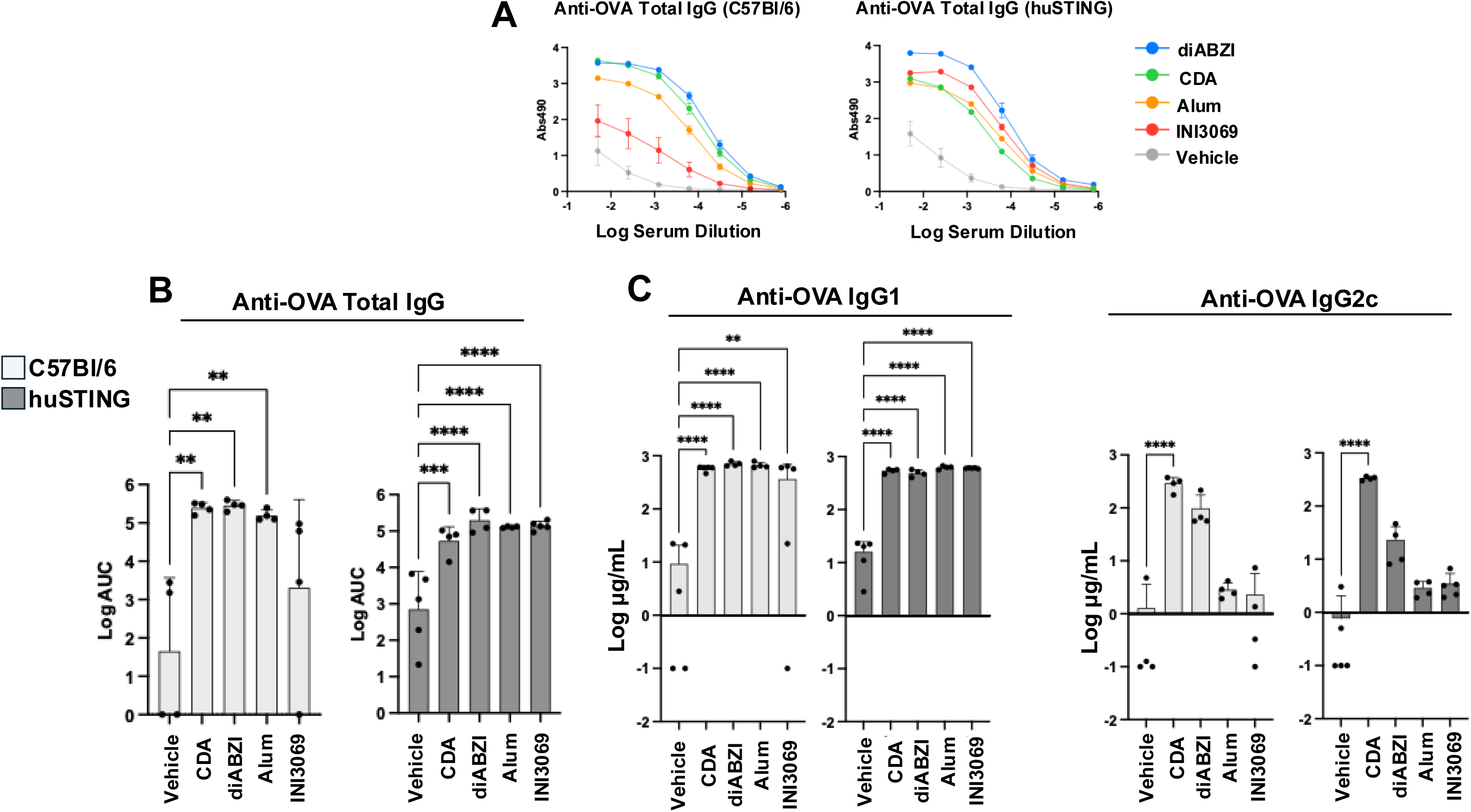
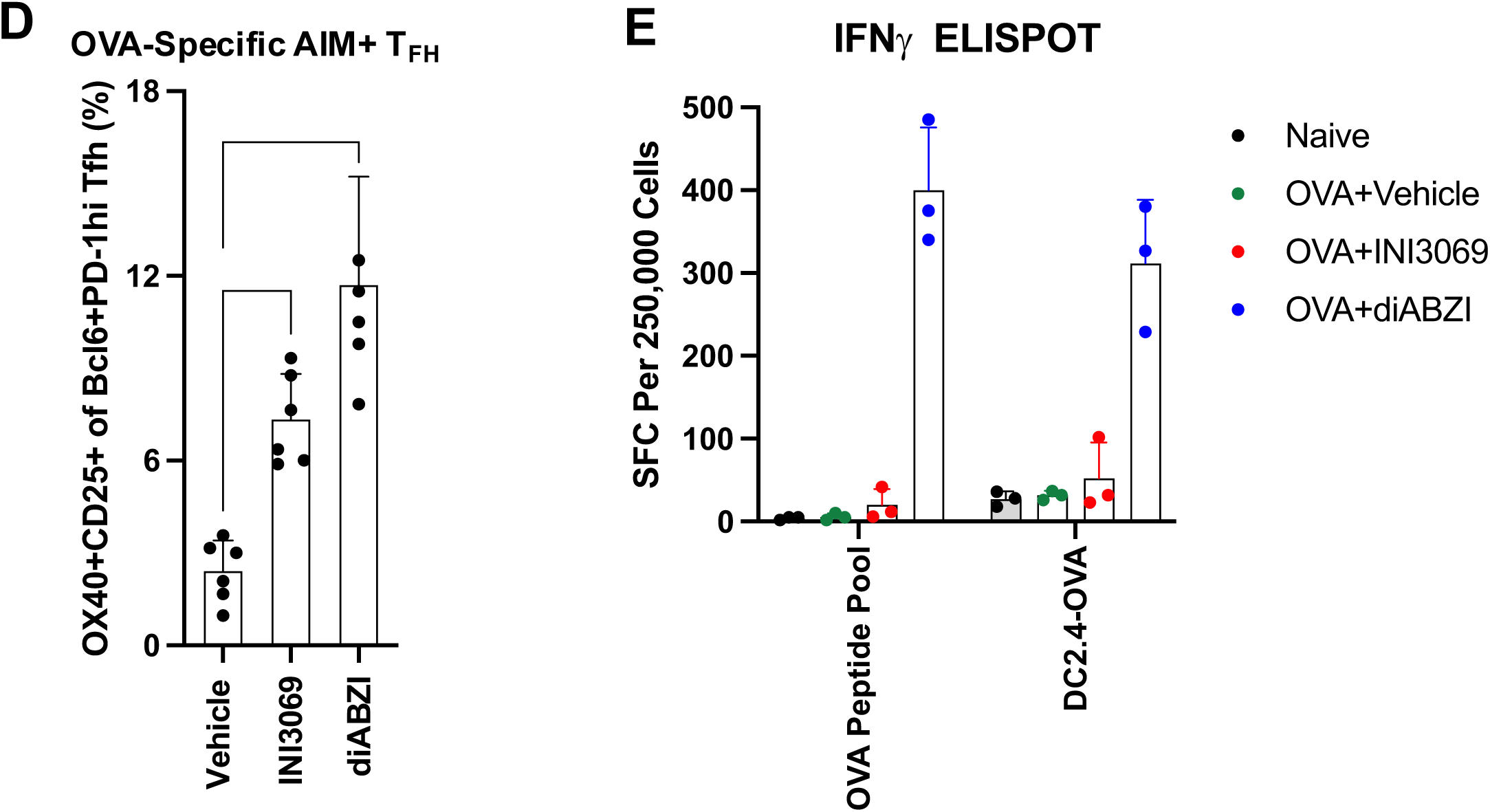
Immune responses to OVA when co-administered with INI3069. C57Bl/6 or huSTING mice (n = 4) were vaccinated SQ with 5 µg OVA in the presence of DMA:PEG vehicle, 10 µg CDA, 10 µg diABZI, 300 µg Alum, or 100 µg formulated INI3069, identically boosted 2 w later, and harvested 2 w later. **A.** Absorbance of serum IgG reactive to OVA as determined by ELISA. Data presented are mean + SEM raw 490 absorbance over indicated serum dilution; **B.** Total IgG titers as determined by area under the curve (AUC) calculations for each adjuvant. Data presented are mean + SEM AUC including individual animal measurements; **C.** Mean + SD OVA-reactive absolute IgG1 and IgG2c levels; **D.** Number of OVA-reactive T_FH_ (CD4^+^CXCR5^+^PD-1^+^) splenocytes harvested seven days after boost with OVA + indicated adjuvant. Statistical significance was determined using ANOVA with Dunnet’s correction for multiple comparisons (*p < 0.05; **p < 0.01; ***p < 0.001; ****p < 0.0001).

Class switch recombination and affinity maturation of antibodies takes place in the lymph node germinal centers and are facilitated by T follicular helper (T_FH_) cells in antigen- and adjuvant-specific manners. We next examined the capacity of STING agonists to induce antigen-associated T_FH_ development in huSTING mice using an activation-induced marker (AIM) assay (89, 90). Animals vaccinated with OVA plus formulation vehicle, diABZI, or INI3069 as above were euthanized, splenocyte single cell suspensions cultured *ex vivo*, and exposed to a peptide pool that spans the OVA protein. Flow cytometry was then used to quantify expression of OX40, CD25, Bcl6, and PD-1 on CD3/4+ cells. As shown in **Figure 10D**, inclusion of diABZI and INI3069 led to a significant increase in the numbers of OVA-reactive T_FH_ cells relative to vehicle alone. Based on these results we conclude that the humoral response in huSTING mice as indicated by antibody formation and antigen-specific T_FH_ activation can be induced by human-selective, TMD-engaging, and LBD-engaging STING agonists.

Next, we examined whether INI3069 can enhance generation of antigen-reactive T cells. For this OVA was coadministered with vehicle alone, diABZI, or formulated INI3069 as above. Splenocytes from these mice were either directly stimulatied with an OVA-spanning peptide pool or mixed with DC2.4 cells stably transduced with full length OVA to skew toward direct antigen presentation. As shown in **Figure 10E**, pairing diABZI with OVA led to antigen specific expression of IFNγ in splenocytes from these mice. However, the use of INI3069 did not (**Figure 10E**). Collectively, these results suggest that INI3069 can enhance antigen-directed IgG1 humoral responses and T_FH_ activation but not IgG2c or cell mediated responses, in contrast to the LBD engaging agonist diABZI. As such, despite the abilities of INI3069 and diABZI to similarly activate STING-mediated TBK1-dependent processes, the immune responses they elicit appear divergent.

### Anti-tumor activity of INI3069 in huSTING mice

Peritumoral injection of STING agonists can lead to immune-mediated clearance of transplanted tumors in murine models [reviewed in (91)]. Demonstrating that syngeneic tumors grow similarly in huSTING and C57Bl/6 mice is essential to establish these animals as suitable for investigating STING-mediated anti-cancer therapies. We therefore subcutaneously injected B16 melanoma and MC38 colon adenocarcinoma cells into C57Bl/6 and huSTING mice and monitored tumor volumes through time. As shown in **Figure 11A** growth rates of B16 and MC38 tumors did not differ significantly between C57Bl/6 and huSTING mice suggesting that the mice should be appropriate for examining anti-tumor treatments.

**Figure 11.**
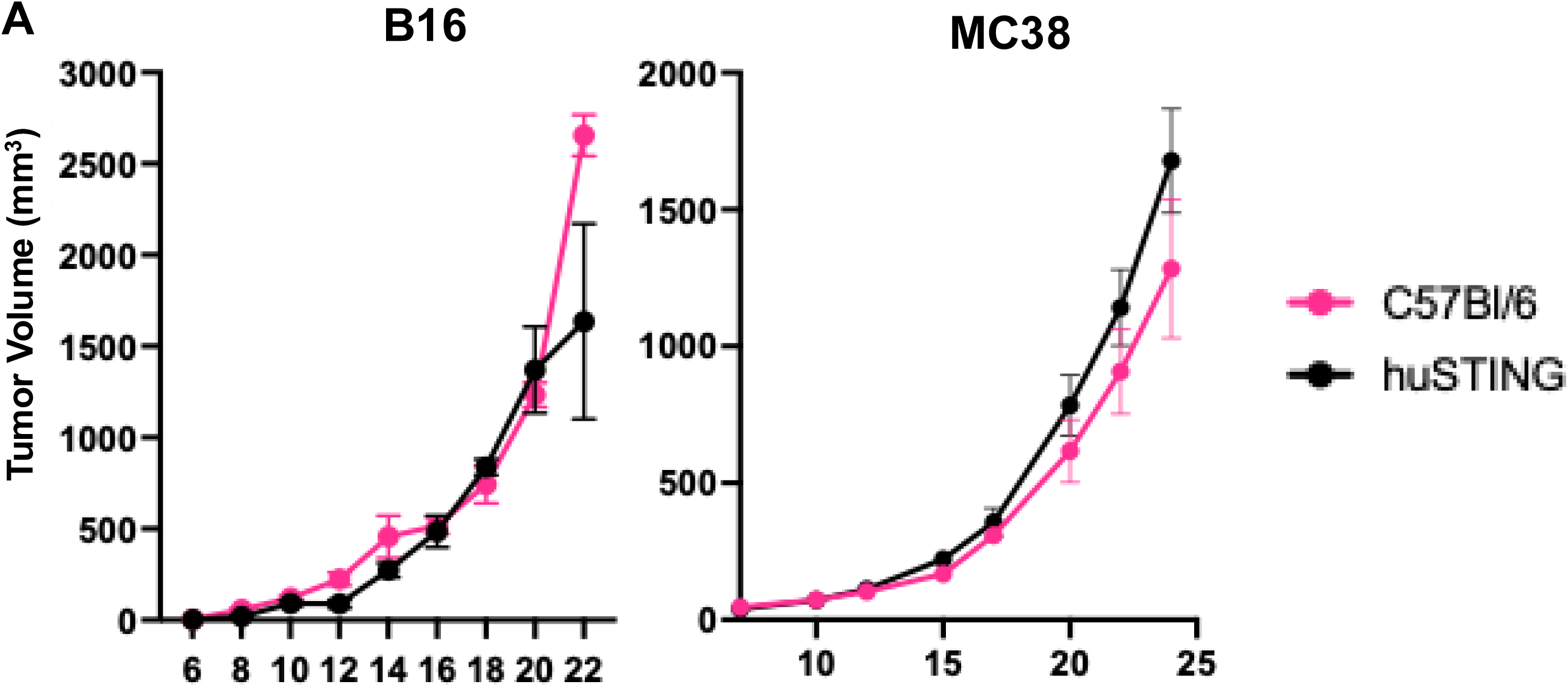
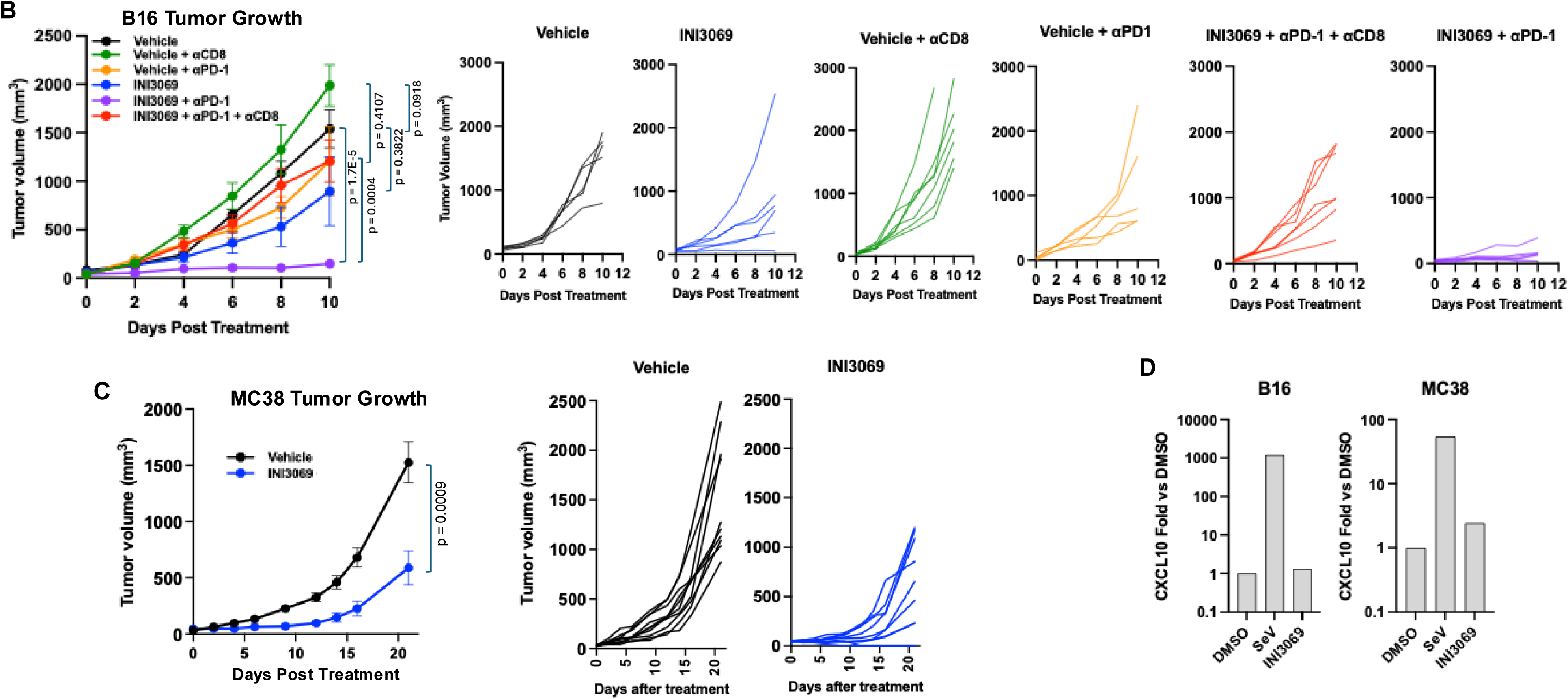
Growth of B16F10 and MC38 Tumors in huSTING Mice Following Treatment with INI3069. **A.** C57Bl/6 and huSTING mice were transplanted SQ with 5 x 10^5^ B16F10 or MC38 cells and tumor volume measured temporally. Data presented are mean ± SEM volumes for indicated timepoint; **B.** huSTING mice were inoculated with 5 x 10^5^ MC38 cells SQ and peritumorally injected three times, 2 days apart with DMA:PEG vehicle or 100 µg formulated INI3069 and tumor size measured. Data presented are mean ± SEM tumor volumes on indicated days post treatment (left) as well as measurements from individual animals (right); **C.** huSTING mice were inoculated with 5 x 10^5^ B16F10 cells SQ and peritumorally injected three times, 2 days apart with DMA:PEG vehicle in the presence or absence of antibody-mediated CD8+ T cell depletion or a-PD-1. 100 µg formulated INI3069 was similarly injected in the presence or absence of a-PD-1 or a-PD-1 during CD8+ T cell depletion. Statistical significance of endpoint tumor volumes was determined using Student’s T-test; **D.** Transcriptional induction of CXCL10 in B16 (left) or MC38 (right) cells following 6 h *in vitro* exposure to SeV (1000 HAU) or 50 µM INI3069. Data presented are mRNA fold change relative to untreated cells as determined by qPCR.

We next asked whether direct administration of INI3069 could impair tumor growth in huSTING mice. For this we first examined B16 cells that are largely insensitive to anti-PD-1 based immune checkpoint inhibition (ICI) therapies (92–94). Transplanted tumors were peritumorally treated three times with formulation vehicle alone or formulated INI3069 and tumor volume monitored (**Figure 11B**). In this approach INI3069 alone did not significantly diminish tumor growth. We next examined whether INI3069 could enhance tumor clearance during ICI by co-administering anti-PD-1 or isotype control antibodies. As shown in **Figure 11B**, this led to profound impairment of tumor growth. Whether CD8^+^ T cells play a role in this outcome was next examined by depleting these prior to and during treatment with INI3069 and anti-PD-1. As shown in **Figure 11B**, depletion of CD8+ T cells alone led to more rapid growth of untreated tumors. Moreover, CD8^+^ T cells were essential for the synergistic INI3069 plus anti-PD-1 effect as tumor growth was largely restored during this treatment scheme in their absence.

To determine whether INI3069 conferred similar effects on the growth of an unrelated tumor type we examined MC38 cells as a model that is more sensitive to ICI. For this we administered only formulation vehicle alone or formulated INI3069 as above. As shown in **Figure 11C**, INI3069 alone led to a significant impairment of MC38 tumor growth. Based on these results we conclude that huSTING mice are a suitable model for examining the anti-tumor responses of human selective STING agonists. Moreover, that INI3069 is capable of impairing growth of an ICI-insensitive tumor in a manner dependent on CD8^+^ T cells as well as impairing the growth of an ICI sensitive tumor. Importantly, INI3069-mediated effects are unlikely to be conferred through a function involving innate activation of the transplanted tumor cells as these are not transcriptionally responsive to the compound as illustrated in **Figure 11D**.

## Discussion

Transient innate induction by STING agonists can generate useful outcomes in multiple clinically important areas including antimicrobial vaccine enhancement (37), antiviral responses (63), antitumor responses (95), and pain suppression (96). While clinical trials involving STING agonists are both planned and ongoing, thus far most have failed and no STING-targeting drugs have received FDA approval (46). The reasons for this are perhaps unique to each entity but beyond pharmacokinetic issues, two factors may contribute to the difference in interspecies performance of STING agonists. First, the evolutionary divergence of STING proteins likely results in ortholog- and allele-specific ligand activity that leads to dissimilar downstream effects. Second, the wide range of underexamined STING-mediated cellular processes means different agonists trigger divergent yet immunologically impactful activities dependent on their chemical structure, region of protein binding, or STING conformational change. Surprisingly, the preclinical criteria used to advance candidate compounds have involved limited molecular phenotypic evaluation of them and conventionally rely on induction of IFN-I as a readout. Yet it is likely that engagement of STING orthologs by specific agonists elicit aggregate response profiles that differ in ways that are not examined. How these processes differ in relation to agonist structure and ortholog has been minimally explored.

We describe potent new analogs of G10 (53) that activate canonical signaling in human cells as indicated by phosphorylation of STING, TBK1 and IRF3 and transcription of ISGs in the presence but not absence of STING. Moreover, they induce activity by three major STING allelic forms and stimulate an IRF3- but not ATG5-dependent antiviral response. Region of STING engagement by agonists can vary and this is functionally significant since activation through the LBD or TMD can differentially induce molecular phenotypes including NLRP3 inflammasome activation, lysosomal biogenesis, and autophagy (23, 24, 47, 69). We used forward genetic and thermal shift methods to demonstrate that INI3069 does not bind to the STING cytosolic domain but requires amino acids in the TMD to induce signaling. This result is consistent with its failure to induce TFEB dephosphorylation and autophagy, responses linked with prevention of STING-mediated proton leakage by TMD obstruction. While we expect INI3069 to display the full spectrum of phenotypes associated with proton blockade this will require additional confirmation.

INI3069 and C53 represent the only known STING agonists to activate the protein by binding to the TMD. However, as human selective compounds, investigation of their *in vivo* effects is not standardly practical. Genetically humanized STING mice, while described previously (97), have been minimally characterized with respect to their cellular and immunological responses and general utility as a clinical model. Cells harvested from huSTING mice respond to cross-reactive (cGAMP, diABZI) and human selective agonists (INI3069 and C53) as indicated by expected activation of phosphorylation cascades, ISG induction, APC maturation, and TFEB dephosphorylation. Based on this we conclude that transgenic human STING recapitulates TBK1-dependent and -independent processes in mouse cells. While an exhaustive assessment of all known STING-mediated processes in these cells was beyond our scope, the investigated phenotypes are known to crucially affect immune activity and associate with both the canonical TBK1-associated and noncanonical proton channel capacities of the protein. Additionally, the primary downstream immunological effects also appear normal in these animals during homeostatic and exogenously activated states. Based on these results we predict that these animals are both uniquely suitable for investigation of human selective STING agonists generally and for characterizing the differential effects of LBD versus TMD engaging compounds specifically.

Subunit vaccine experiments indicate that the animals respond to exogenous STING induction in a manner that resembles what is observed in wild type mice and as such are appropriate for these studies. Furthermore, they reveal qualitative differences in immunogenicity by INI3069 and LBD-binding agonists such as CDA and diABZI. For instance, a T_H_2 biased response is observed for INI3069 whereas more balanced T_H_1/T_H_2 polarization is seen for LBD agonists as indicated by relative levels of IgG1 and IgG2c (**Figure 9A-C**). Additionally supportive of this is the finding that diABZI, but not INI3069, is capable of eliciting cell mediated responses to OVA as indicated by IFNγ^+^ ELISPOT (**Figure 9E**). However, both compounds can stimulate T_FH_ differentiation, which is consistent with high IgG1 antibody responses following INI3069 administration (98) (**Figure 9D**). Whether these results are mechanistically linked with differential molecular processes induced by LBD and TMD agonists is a crucial question with implications for STING agonist development. For instance, TFEB is emerging as playing a major role in adaptive immunity including guiding presentation by DC of exogenous antigens toward MHC-II and away from MHC-I (72), expression of CD40 ligand critical for T and B cell interactions and antibody production (99), and regulatory T cell differentiation (100). As such, the observation that, unlike LBD agonists, TMD agonists such as INI3069 fail to activate TFEB (**Figures 4B** and **8B**) suggests that STING agonist classes can lead to immunologically distinct outcomes.

The enhancement of innate and antigen-directed immune responses by INI3069 is suggestive of its potential for display of anti-tumor effects as has been shown for other STING agonists. We examined this by directly treating transplanted MC38 and B16 tumors, which grow at typical rates in huSTING mice (**Figure 10A**). These tumor types were selected due to both their clinical relevance and their differential reaction to immune checkpoint inhibition therapies with MC38 being more (101) and B16 less (102) sensitive. These results indicate that when administered alone, INI3069 is able to significantly slow MC38 tumor growth (**Figure 10B**). While INI3069 monotherapy was unable to similarly slow B16 tumor growth, it blocked growth in the presence of anti-PD-1 ICI and this is likely mediated by CD8^+^ T cells since their depletion abrogated this effect (**Figure 10C**). Future studies will be needed to establish the degree to which INI3069 anti-tumor properties are mediated by antigen-based or proinflammatory processes. However, given the human selectivity of the molecule and its demonstrated activity on human cells (**Figure 3**), it represents an unusually promising clinical candidate for further development.

*In vivo* activity of any kind has not been previously demonstrated for TMD-engaging STING agonists. As such this work shows that this class of molecule can be effective at stimulating translationally relevant outcomes. Furthermore, these results illustrate that in addition to the demonstrably different molecular processes triggered by TMD and LBD engaging molecules (23, 24, 69), differences in T helper polarization are also evident. Our group has recently shown that chemically diverse LBD-engaging STING agonists stimulate transcriptomic, molecular, innate, and adaptive immune responses that can diverge substantially between molecules (103). We therefore expect that characterizing the totality of such phenotypes induced by INI3069 and C53 will be both necessary for their clinical advancement and reveal novel phenomena about the physiologic roles of STING. Fortunately, this work demonstrates that huSTING mice are valuable for this pursuit in the context of immunological processes that will allow preclinical inquiries into vaccine responsiveness and anti-tumor effects.

## Materials and Methods

### Reagents and antibodies

diABZI was purchased from MedChemExpress (#HY-112921B). DMXAA was purchased from ApexBio (# A8233). Sendai virus (SeV) was purchased from Charles River (# PI-1). Steady GLO (# E25100), Nano-Glo (# N1120), and Cell Titer Glo (# G7572) were purchased from Promega. Quanti-Luc (# rep-qlc4lg1), endotoxin-free ovalbumin (# vac-pova-100), alhydrogel aluminum hydroxide (# vac-alu-250), 2’3’ cGAMP (# tlrl-nacga23-02), CDA derivative ML-RR-S2-CDA (# tlrl-nacda2r-01), were purchased from Invivogen. Lipofetamine 3000 was purchased from ThermoFisher (# L3000015). LPS was purchased from Sigma-Aldrich (# L5293). N,N-dimethylacetamide anhydrous (DMA; # 271012) and polyethylene glycol 400 (PEG; # 91893) were purchased from Sigma-Aldrich. Anti-mouse β-Actin (# 5125), anti-mouse/human STING (# 13647), anti-mouse phospho-S366 STING (# 72971), anti-human phospho-S365 STING (# 19781), anti-mouse/human IRF3 (# 4302), anti-mouse phospho-S379 IRF3 (# 79945), anti-mouse/human TBK1 (# 3504), anti-mouse/human phospho-S172 TBK1 (# 5483), and anti-phospho-S211 TFEB (# 37681S) were purchased from Cell Signaling. Anti-mouse CD3e (# 565992), anti-mouse Bcl6 (# 563363), anti-mouse CD25 (# 564458), anti-mouse MHC-II (# 553823) were purchased from BD Biosciences. Anti-mouse CD4 (# 100447), anti-mouse CD45R/B220 (# 103243), anti-mouse PD-1 (# 135205), anti-mouse OX40 (# 119413), anti-mouse CD40 (# 124622), anti-mouse CD80 (# 124622), anti-mouse CD86 (# 105008), and anti-mouse CD11c (# 117353) were purchased from BioLegend. Anti-mouse PD-1 (# BE0146), anti-mouse CD8α (# BE0061) were purchased from Bio-X-Cell. Anti-TFEB was purchased from Bethyl (# A303-673A-T). Anti-mouse IgG-HRP (# 1030-05), anti-mouse IgG1-HRP (# 1070-05), and anti-mouse IgG2c-HRP (# 1079-05) were purchased from Southern Biotech.

### Luciferase reporter assays

THF reporter or HEK293T cells were plated at 20,000 cells per well in a white 96-well plate 24 h before stimulation. Treatments were performed in duplicate overnight in 50 µL DMEM plus 2% FCS. One-GLO lysis/luciferin reagent was added 1:1 to each well, and luminescence measured on a Bio-Tek Synergy plate reader. Reporter assays using HEK293T cells stably transduced with STING alleles involved plating 500,000 cells in 35 mm dishes and transfecting plasmids encoding IFIT1-dependent luciferase (IFIT1-Luc) using Lipofetamine 3000 according to manufacturer’s instructions. After 24 h cells were trypsinized and plated at 20,000/well in 96 well plates and exposed in duplicate to indicated stimuli eluted in 50 µL DMEM plus 2% FCS for 24 h and luminescence read as above.

### Immunoblotting

Sodium dodecyl sulfate-polyacrylamide gel electrophoresis (SDS-PAGE) immunoblotting was performed as follows. After trypsinization and cell pelleting at 2,000 x g for 10 min, whole-cell lysates were harvested in RIPA lysis buffer (50 mM Tris-HCl [pH 8.0], 150 mM NaCl, 1% NP-40, 0.5% sodium deoxycholate, and 0.1% SDS) supplemented with protease and phosphatase inhibitor cocktail. Lysates were electrophoresed in 8% polyacrylamide gels and transferred onto polyvinylidene difluoride membranes using semidry transfer at 400 mA for 1 h. The blots were blocked at room temperature for 2 h or overnight using 10% nonfat milk in 1 x PBS containing 0.1% Tween 20. The blots were exposed to primary antibody in 5% nonfat milk in PBS containing 0.1% Tween 20 for 18 h at 4°C. The blots were then washed in PBS containing 0.1% Tween 20 for 20, 15, and 5 min, followed by deionized water for 5 min. A 1 h exposure to horseradish peroxidase-conjugated secondary antibodies and subsequent washes were performed as described for the primary antibodies. The antibody was visualized using enhanced chemiluminescence.

### Construction of stable cell lines and genome editing

Stable overexpression of STING alleles and mCherry-GFP-LC3 fusion protein in HEK293T cells and OVA in DC2.4 cells or CRISPR/Cas9-mediated gene deletion of THF cells was performed using lentiviral transduction. DNA encoding WT, HAQ, and R232H STING alleles (using identical codon optimization patterns), mCherry-GFP-LC3, or OVA sequences were synthesized by Twist Biosciences. These were cloned into the pLVX-EF1a-Puro vector (Takara Bio). For CRISPR/Cas9-mediated knockout of ATG5, targeting gRNA (GATGGACAGTTGCACACACT) was cloned into LentiCRISPRv2 (AddGene #98290) as described (104). Lentivirus for these constructs was made by transfecting them along with pMD2.g (AddGene #12259) and psPAX2 (AddGene #12260) into LentiX cells (Takara Bio) using Lipofectamine 3000. At 24 h media was harvested, cleared by centrifugation, and filtered through 0.45 µm filter. This was then added to target cells in the presence of 8 µg/mL polybrene. Transduced cells were selected by exposing to 1 µg/mL puromycin. Single cell clones were then obtained either by serial dilution or fluorescence activated cell sorting (mCherry-GFP-LC3) and target protein expression or deletion verified using immunoblot.

### Ratiometric flow cytometry

HEK293T cell clones stably transduced with mCherry-GFP-LC3 were either left untrasfected or transiently transfected for 24 h with human pLVX-EF1α plasmid encoding STING (WT). Cells were exposed to 1 µM diABZI, 10 µM C53, or 10 µM INI3069 for 4 h. Flow cytometry was then performed using A5 Symphony 28 (BD Biosciences) and analyzed using Flowjo v10.10.0 (Tree Star). The ratio of MFI for mCherry to GFP was determined for individual cells. Degree of autophagic flux was assigned based on the distribution of ratios observed in cell populations as shown in **Figure 5B** and as described in (105, 106). Experiments were performed in two independent clonal cell lines in duplicate. A replicate experiment was also performed and whole cell lysates harvested from these for immunoblot.

### In vitro virus infection

Stocks of Nanoluciferase expressing CHIKV (CHIKV-nLuc) was derived from an infectious clone as previously described (65). Live virus was generated from *in vitro* transcribed RNA that was then transfected via Lipofectamine into BHK-21 cells and media collected. Resultant virus was titered as above and propagated in C6/36 insect cells for 48 h to produce viral stocks. Virus was pelleted through a 20% sucrose cushion by ultracentrifugation (22,000 rpm [825,206 x g] for 1.5 h). Experimental infections of indicated cells were carried out in triplicate using a multiplicity of infection (MOI) of 0.1 PFU per cell. CHIKV-nLuc was quantified by adding Nano-Glo luciferin lysis buffer (Promega) to cells at a 1:1 ratio and reading luminescence as described above.

### Hybridization array transcriptomic analysis

Microarray assays were performed in the OHSU Integrated Genomics Laboratory. RNA sample quantity and purity were measured by UV absorbance with a NanoDrop 1000 spectrophotometer. RNA integrity and size distribution were determined by running total RNA on a Nano chip instrument (Agilent Technologies). RNA was prepared for array hybridization by labeling 100 ng aliquots using the 3’IVT Express kit (Affymetrix). RNA was reverse transcribed to generate first-strand cDNA containing a T7 promoter sequence. A second-strand cDNA synthesis step was performed that converted the single-stranded cDNA into a dsDNA template for transcription. Amplified and biotin-labeled cRNA was generated during the in vitro transcription step. After a magnetic bead purification step the cRNA was fragmented. Labeled and fragmented cRNA was combined with hybridization cocktail components and hybridization controls, and 130 µL of each hybridization cocktail containing 6.5 µg of labeled target was injected into a cartridge containing the Clariom S murine array (Affymetrix) interrogating over 150,000 transcripts from >22,100 genes. Arrays were incubated for 18 h at 45°C, followed by washing and staining on a GeneChip Fluidics Station 450 (Affymetrix) and the associated hybridization wash and stain kit. Arrays were scanned using the GeneChip Scanner 3000 7G. Image processing of sample .DAT files to generate probe intensity .CEL files was performed using the Affymetrix GeneChip Command Console (AGCC) software. Each array file was then analyzed using Transcriptome Analysis Console (TAC; Version 4.0.3) to obtain array performance metrics and calculate transcript fold changes. To identify probe sets that were significantly regulated in treated versus untreated (mock) cells, we employed a traditional unpaired one-way (single-factor) ANOVA for each pair of condition groups as implemented in TAC. Probe sets were considered differentially regulated if the ANOVA P value was <0.05 and fold change >2. Pathway analysis of induced transcripts was performed using Ingenuity Pathway Analysis (Qiagen).

### Construction of transgenic knock in mice

Genetically humanized STING mice were constructed by Cyagen Biomodels, LLC. Briefly, the sequence encompassing exon 3 to exon 8 of mouse Sting1 was replaced by the Human STING1 CDS-polyA-sdNeo cassette shown in **Figure 6**. The targeting vector was electroporated into C57BL/6N embryonic stem (ES) cells. Cells were then cultured in a medium containing neomycin. PCR genotyping was then used to assess acquisition of the desired mutation through homologous recombination. Targeted ES cell clone 1H2 was injected into C57BL/6 embryos, which were then re-implanted into CD-1 pseudo-pregnant female chimeras. The resulting progeny (F0) were then bred with wildtype animals and selected for germline transmission. Among the next generation (F1), the globe humanized mice were verified by PCR screening. The heterozygous F1 were further intercrossed for homologous mutant F2. These mice express the human STING1 allele under the control of endogenous mouse Sting1 promoter with no mouse Sting1 expressed.

### In vivo *studies*

C57Bl/6 were purchased from the Jackson Laboratory. huSTING mice were bred in house. All animals were used at 6 - 8 weeks of age. For vaccination experiments animals were injected subcutaneously with 5 µg endotoxin free OVA admixed with 200 µg INI3069 formulated in DMA:PEG, 10 µg diABZI, 10 µg ML-RR-S2 (CDA), 300 µg Alhydrogel, or DMA:PEG vehicle alone in 100 µL total volume and boosted identically at 2 w. For INI3069 formulation a 20 mg/mL stock in DMA was combined with 10 volume parts DMA was combined with 30 volume parts PEG, and 60 volume parts PBS. For serum antibody studies animals were euthanized at 2 w post boost and for ELISPOT studies animals were euthanized at 1 w post boost. For tumor studies huSTING mice were injected subcutaneously (SQ) with 5 x 10^5^ B16F10 or MC38 cells and tumor volume (V) monitored every other day using calipers (V = 0.5 x L x W^2^). When tumor sizes reached 100 mm^3^ three peritumoral injections of vehicle alone or 100 µg formulated INI3069 were performed every other day. Immune checkpoint inhibition was induced by intraperitumorally (IP) injected 200 µg anti-PD1 antibody three times every other day at the start of tumor treatment. CD8+ T cell depletion was performed by IP injection of 100 µg anti-CD8a antibody every four days starting two days before tumor inoculation. CD8+ cell depletion was validated by flow cytometry performed on peripheral blood collected by submandibular bleed.

### RNA isolation and quantitative reverse transcription-PCR

Total RNA was isolated from cells or tissues, treated with DNase provided in a DNA-free RNA isolation kit (Zymo Research) according to the manufacturer’s protocol, and quantified using UV spectrometry. Single-stranded cDNA for use as a PCR template was made from total RNA and random hexamers to prime first-strand synthesis via reverse transcriptase. Relative fold change of mRNA expression between treated and mock-treated samples was calculated using semi-quantitative real-time RT-PCR (qPCR) with the Applied Biosystems sequence detection system according to the ΔΔCT method (107) with GAPDH as a control. mRNA-specific pre-validated Prime-Time 6-carboxyfluorescein qPCR primer/probe sets obtained from IDT were used for all genes. Absolute quantitation of viral DNA copies or mRNA transcripts from mouse tissues was performed by isolating RNA and synthesizing cDNA as above. A serial dilution of plasmid encoding the virus-specific target region was included to establish a Ct value standard curve which allowed regression analysis to establish absolute mRNA copy numbers per µg total RNA.

### STING thermal shift assays

Engagement of purified human STING C-terminal domain and INI3069 was performed as described (108). Briefly, an open reading frame encoding 6xHIS tagged STING amino acids 137 to 379 was cloned into pRSET-B and used to transform the *E. coli* strain pLysS. Liquid cultures of transformed *E. coli* were then induced to express the protein using 1 mM isopropyl-D-thiogalactopyranoside at 16°C for 18 h. STING protein was purified by nickel-affinity chromatography (Clontech Laboratories) and purified by gel filtration chromatography (HiPrep 16/60Sephacryl S-100 HR column; GE Healthcare Life Sciences). Eluted proteins were concentrated using Amicon centrifugal filters (10-kDa cutoff). For thermal shift assay, 1 μg of recombinant protein was combined 100 µM 2’3’-cGAMP or 50 µM INI3069 along SYPRO Orange dye at 1:1000 dilution in a 20μL reaction (in triplicate). A StepOne Plus Real-time PCR system was used to acquire fluorescence over a temperature gradient of 25 to 99°C. The melting curves were plotted as fluorescence signal using GraphPad Prism 10 software.

### MEF isolation, bone marrow cell harvest, and DC maturation

Murine embryonic fibroblasts (MEF) were harvested from C57Bl/6, huSTING, and C57Bl/6^Gt/Gt^ and cultured as described (109). Immature bone marrow derived dendritic cells (BMDC) were obtained by collecting marrow from mouse femurs using syringe flushing followed by red blood cell lysis in ACK buffer. Cells were cultured in RPMI + 10% FCS in the presence of 20 ng/mL GM-CSF and 10 ng/mL IL-4 (Peprotech). Cytokines and media were replaced on day 3. Nonadherent cells were removed and media similarly replaced on day 6. On day 8 cells were then exposed to indicated stimuli for 24 h and flow cytometry performed. Agonist stimulation was performed for 24 h with 0.5% DMSO, 10 nM diABZI, or 10 µM INI3069.

### BMDC flow cytometry

Cells were collected by centrifugation at 500 x g for 5 minutes and washed with FACS buffer (0.5% BSA, 5 mM EDTA in PBS). Cells were blocked with 5 µg/mL FC block and dead cells were stained with Live/Dead Fixable Near IR at 4°C for 15 minutes. Staining antibodies CD11c, MHC-II, CD80, CD86, and CD40 were diluted and incubated with cells at 4°C for 30 minutes and washed. Data acquisition was performed using A5 Symphony 28 (BD Biosciences) and analyzed using Flowjo v10.10.0 (Tree Star). Median MFI of activation markers were compared to DMSO control.

### Activation induced marker (AIM) assay

Splenocytes from vaccinated mice were incubated with 2 µg/mL 15-mer OVA peptide pool (Miltenyi) at 37◦C with 5% CO_2_ for 18h. Stimulation with 5 µg/mL concanamycin A was included as a positive control. After stimulation, cells were stained with LIVE/DEAD Fixable Near IR and blocked with mouse Fc block CD16/CD32 (BD Biosciences) at 4°C for 15 minutes. Cells were then stained for 30 min at 4°C with the following fluorochrome-conjugated anti-mouse Abs: CD3, CD4, CD45R/B220, CD25, PD-1, and OX40. Intracellular staining was performed for 45 min at 4◦C with anti-mouse Bcl6 antibody using Cytofix/Cytoperm Fixation/Permeabilization kit (BD Biosciences), according to manufacturer’s instructions. Cell events were collected on A5 Symphony 28 (BD Biosciences) flow cytometer, and data were analyzed using FlowJo v10.10.0 software.

### IFNγ ELISPot

Splenocytes were harvested from mice one week after vaccination boost and maintained in prewashed mouse IFNγ ELISPot plates (MabTech) at 250,000/well in RPMI + 10% FBS following red blood cell lysis. They were either stimulated with a pool of 15-mer OVA inclusive peptides at 10 µg/well or mixed with DC2.4 cells that stably overexpress full length OVA at a splenocyte:DC2.4 ratio of 1:10 for 18 h. Plates were then stained according to the manufacturer’s protocol. Briefly, plates were washed and incubated with anti-mouse IFNγ biotin antibody for 2 h. Steptavidin-ALP secondary antibody was added for 1 h after plates were washed. Spots were detected using BCIP/NPT-plus substrate, rinsed with water, and dried before counting with an AID ELISpot plate reader.

### Enzyme-linked immunosorbent assay (ELISA)

To measure levels of antigen-specific total IgG in mouse serum sandwich ELISA was used. OVA-reactive antibodies were measured by first coating 96 well plates with OVA eluted in 100 µL PBS at 10 µg/mL at 4°C for 24 h. Plates were blotted dry and blocked for 1 h with 200 μL of 5% milk in PBS with 0.05% Tween-20 (ELISA buffer). Plates were then washed three times with 200 μL of ELISA buffer. Serum was heat-inactivated at 56°C for 0.5 h, diluted 1:50 in ELISA buffer, and serially diluted 1:5 seven times. 100 μL of each dilution was added to wells in duplicate and incubated for 1.5 h at room temperature (RT). Plates were washed three times with ELISA buffer and blotted dry. HRP conjugated secondary antibodies were diluted 1:10,000 in ELISA buffer and 100 μL was added to wells and incubated for 1 h at RT. Plates were then washed three times with ELISA buffer, blotted dry and developed with HRP substrate. The reaction was stopped with 1 M HCl 10 minutes after exposure. Absorbance was then read at 490 nm on a BioTek Synergy plate reader. Total IgG titers were calculated by determining area under the curve for serum-specific absorbance and log transformed. Absolute levels for antigen-reactive IgG1 and IgG2c were determined by using linear regression based on a standard curve of control isotype specific mouse antibodies.

### Ex vivo PBMC collection and stimulation assay

Human peripheral blood mononuclear cells (PBMCs) were isolated from multiple cohorts of healthy adult donor by leukapheresis. Single-cell PBMC suspensions were frozen in bovine serum albumin (MilliporeSigma) plus 10% DMSO for cryopreservation in liquid nitrogen and thawed from for experiments in this study. Thawed PBMCs were transferred to 50 mL polypropylene tubes, spun down and suspended in cultured RPMI medium with 10% FBS and 1X (50U) and penicillin-streptomycin. For stimulation, PBMCs were plated at 10^6^ cells/well in 96-well polypropylene deep well plates. Monocyte-derived immature DCs were generated from PBMCs. Monocytes were enriched from leukapheresed blood using a CD14^+^ monocyte magnetic bead separation kit without CD16 depletion (Stemcell). Enriched monocytes were differentiated into immature DCs using 100 ng/mL GM-CSF (Gemini) and 20 ng/mL IL-4 (Gemini) in CellGenix GMP DC medium (Sartorius), and harvested after differentiating for 36-48 hours at 37°C with 5% CO_2_. For stimulation of immature DCs, cells were plated at 500,000 cells/well in 96-well polypropylene deep well plates. PBMCs or immature DCs were plated with INI3069 (5, 10 µM) or dose matched DMSO overnight at 37°C with 5% CO_2_. Supernatant and cells were harvested for subsequent multiplex immunoassay analysis flow cytometric analysis, respectively.

### Human PBMC flow cytometry analysis

For *ex vivo* analysis of activation and maturation marker expression, 10^6^ PBMCs or immature DCs stimulated with varying concentrations of INI3069, were incubated with Live/Dead Fixable Aqua viability stain 405 (ThermoFisher Scientific) to discriminate dead from live cells then stained with incubated in 1X Annexin-V buffer with fluorochrome-conjugated antibodies for at least 15–20 minutes at 4°C or on ice, protected from light. The following fluorochrome-conjugated anti-human antibodies from Biolegend or BD were used: CD40-APC, PD-L1 PE-CF594, ICOSL-BV421, CD83-PE, CD86-BV605, CD80-PECy5, HLA-DR-AF700, CD14, BV650, CD16-PECy7, Annexin-V-FITC, and a DUMP (CD3, CD19) channel on APC-Cy7. Flow cytometry measurements were made on a BD LSRFortessa (BD Biosciences) and collected data analyzed using FlowJo software version 10.7.1.

### Multiplex cytokine assay

Cytokines/Chemokines were measured in an overnight culture supernatant from PBMC and immature DC stimulation experiments using the Human Chemokine Panel, 40-Plex (Bio-Rad, CA, USA) according to the manufacturer’s instructions. The following human analyte premixed panels were used: 6Ckine / CCL2, BCA-1 / CXCL13, CTACK / CCL27, ENA-78 / CXCL5, Eotaxin / CCL11, Eotaxin-2 / CCL24, Eotaxin-3 / CCL26, Fractalkine / CX3CL1, GCP-2 / CXCL6, GM-CSF, Gro-α / CXCL1, Gro-β /, CXCL2, I-309 / CCL, IFN-L, IL-1β,, IL-2, IL-4, IL-6, IL-8 / CXCL8, IL-10, IL-16, IP-10 / CXCL10, I-TAC / CXCL11, MCP-1 / CCL2, MCP-2 / CCL8, MCP-3 / CCL7, MCP-4 / CCL13, MDC / CCL22, MIF, MIG / CXCL9, MIP-1α / CCL3, MIP-1δ / CCL15, MIP-3α / CCL20, MIP-3β / CCL19, MPIF-1 / CCL23, SCYB16 / CXCL16, SDF-1α+β / CXCL12, TARC / CCL17, TECK / CCL25, TNF-α. Supernatant samples were added 1:2 to the plate and cytokine standards were supplied by the manufacturer. The samples were acquired using a Bioplex-200 system and the data analyzed on the BioPlex Manager Software.

### Study Approval

All mouse experiments were performed at Oregon Health and Science University in ABSL2 laboratories under compliance granted by the Institutional Animal Care and Use Committee (IACUC), under protocol TR01_IP00003648. The OHSU IACUC adheres to NIH Office of Laboratory Animal Welfare standards (OLAW welfare assurance A3304-1). Human samples were obtained at Drexel University College of Medicine (PA, USA), Martin Memorial Health Systems (FL, USA), and the Chronic Viral Illness Service at McGill University Health Centre (Montreal, Quebec) after all participants provided signed informed consent under IRB # 083.

### Data Availability

The raw transcriptomic data this study is available through the Gene Expression Omnibus (https://www.ncbi.nlm.nih.gov/geo/) with the accession number to be determined upon approval.

## Supporting information

Supplemental Table 1 transcriptomics

Supplemental Table 2 Pathway analysis

Summplemental methods

## Author Contributions

- Conceptualization: VD, OR, DB, EH
- Methodology: VD, OR, Dave B, EH, NM, SM
- Experimentation: NM, JA, KJ, IR, LS, Dylan B, TA, DS, JW, SM, UP, AJ, DJ, RM, OR
- Funding acquisition: VD, EH, Dave B
- Project administration: VD, OR, EH
- Supervision: VD
- Writing: VD

