## Supplementary material for "Characterization of a Novel Transmembrane Activating STING Agonist using Genetically Humanized Mice": Summplemental methods

**Chemistry:**

By introducing electron donating groups at the 2 -position of the fluorine-containing aromatic ring of the known STING agonist G10, we have designed a new series of compounds with enriched electron density on the aromatic ring. The chemical synthesis of these STING agonists is shown in Scheme 1. Unless otherwise stated, all starting materials are commercially available. Arylation reaction of methyl 3-oxo-2H-1,4-benzothiazine-6-carboxylate (**1**) with substituted benzyl bromide gave compound (**2**), which underwent a hydrolysis reaction to give the key intermediate (**3**). Compound (**3)** was treated with aryl or furfuryl methyl amine to give the corresponding substituted analogs **INI3067**, **INI3074** and **INI3075** respectively, which were subsequently reacted with BBr_3_ leading to the corresponding products **INI3067**, **INI3074** and **INI3075**.

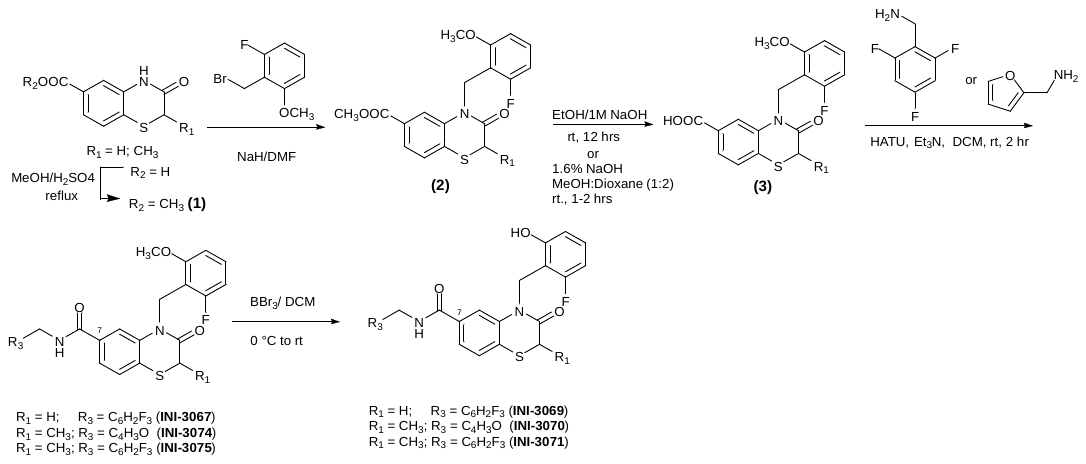

**Scheme 1:** Synthesis of Compounds **INI3069**, **INI3070** and **INI3071**.
